# The genome trilogy of *Anopheles stephensi*, an urban malaria vector, reveals structure of a locus associated with adaptation to environmental heterogeneity

**DOI:** 10.1101/2021.08.25.457468

**Authors:** Aditi Thakare, Chaitali Ghosh, Tejashwini Alalamath, Naveen Kumar, Himani Narang, Saurabh Whadgar, Kiran Paul, Shweta Shrotri, Sampath Kumar, M Soumya, Raksha Rao, Mahul Chakraborty, Bibha Choudhary, Susanta K. Ghosh, Suresh Subramani, Sunita Swain, Subhashini Srinivasan

## Abstract

**Background:** *Anopheles stephensi* is the most menacing malaria vector to watch for in newly urbanizing parts of the world. The fitness is reported to be a direct consequence of the vector adapting to laying eggs in over-head water tanks with street-side water puddles polluted by oil and sewage. Large frequent inversions of malaria vectors are implicated in adaptation.

**Results:** We report the assembly of a strain of *An. stephensi* of the type-form, collected from a construction site from Chennai (IndCh) in 2016. The genome completes the trilogy with respect to a 16 Mbp inversion (2R*b*) in *An. stephensi* associated with adaptation to environmental heterogeneity. Comparative genome analysis revealed breakpoint structure and allowed extraction of 22,650 segregating SNPs for typing this inversion. Using whole genome sequencing of 82 individual mosquitoes, we conclude that one third of both wild and laboratory populations maintain heterozygous genotype of 2R*b*. The large number of SNPs are tailored to assign inversion genotype directly from 1740 exonic SNPs 80% of which are expressed in various developmental stages.

**Conclusions:** The genome trilogy approach accelerates study of fine structure and typing of important inversions in malaria vectors putting the genome resources for the much understudied *An. stephensi*, on par with the extensively studied malaria vector, *Anopheles gambiae*. We argue that the IndCh genome is relevant for field translation work compared to those reported earlier by showing that individuals from diverse populations cluster with IndCh pointing to significant commerce between cities, perhaps, allowing for survival of the fittest strain.

## BACKGROUND

*Anopheles (An.) stephensi* is a malaria vector prevalent in urban India, South Asia, Middle East and, more recently, expanding its range into Africa as urbanization continues[1]. The persistence of *An. stephensi* in urban settings is linked to its ability to lay eggs in overhead water tanks, when the stagnant water on the streets is rendered unsuitable[2]. The genomes of a few strains of *An. stephensi* have been reported recently, including that of a laboratory strain of *An. stephensi*, STE2, MRA-128, originally collected from Delhi by the NIMR and subsequently maintained at the Walter Reed Hospital[3]. The STE2 strain has been extensively studied, leading to several publications including the complete physical map[4] and a chromosomal study of several inversions via photomaps of loop formation, including the heterozygous 2R*b* inversion[5]. As early as 2014, using a cocktail of the then state-of-the-art sequencing technologies, this group sequenced DNA extracted from a pool of 50 lab-reared individuals to assemble a draft genome[6]. Several recent attempts improve this draft genome using synteny[7], homology-based approaches[3] and paired-end HiC reads[8] to obtain a near-chromosome level assembly, which confirms the presence of the 2R*b* inversion in heterozygous form (2R+^b^/2R*b*)[3]. This report catalogued the majority of the coding genes and their gene structures[6] in *An. stephensi,* for the first time.

More recently, a very high-resolution genome was reported for a laboratory strain, hereafter referred to as the UCI strain, providing a gold-standard reference genome in guiding future malaria research[9]. The contig-level statistics of this assembly are close to those of the human and fly genomes, the two that are the most complete to date. Based on this assembly and reconciliation with HiC data, it has been reported that the UCI genome is homozygous for the 2R*b* inverted form (2R+^b^/2R+^b^)[3].

The importance of paracentric inversions in *An. gambiae* has been established and proposed to be associated with the ability of some vectors to follow humans in diverse environments[10]. By studying polytene chromosomes from 1500 individuals of *An. gambiae* from the forest and the savannas, it was reported that those from the forest are homozygous for standard forms, but those from the savannas display complex inversion patterns in the chromosome 2 of *An. gambiae*. Of these, the 2R*b* and 2L*a* inversions are the most well-studied in terms of their frequencies associated with traits[11][12]. It was proposed that some inversions may be as old as the expansion of agriculture and may have been maintained in the population for a long time giving much time for genetic drift between the heterozygous arms.

As early as 1972, polytene chromosomes from the ovarian nurse cell in adult *An. stephensi* mosquitoes have been used to study the association of inversion polymorphisms with certain experimental or ecological situations[13]. At that time, the 2R*b* inversion was known as the Karachi variant (KR). Using *An. stephensi* strains from four distal geographical locations including Iraq, Iran, Pakistan and India, this group reported that the strains from India were homozygous for the standard form of the 2R*b* configuration (2R+^b^/2R+^b^); those from Iraq/Iran were heterozygous and that from Karachi was homozygous for the 2R*b* inversion (2R*b*/2R*b*). The inversion karyotype was correlated with early emergence for homozygous 2R+^b^/2R+^b^, intermediate emergence for heterozygous form 2R+^b^/2R*b* and late emergence for homozygous 2R*b*/2R*b* form. Also, it was reported that a cross between opposite homozygous forms did not produce intermediate emergence, suggesting a selection process that is more complex. They studied the feeding behavior of 2R+^b^/2R+^b^, 2R+^b^/2R*b* and 2R*b*/2R*b* genotypes of Iraq strains and noticed differences in sugar-feeding propensity during photophase, implicating this locus in the control of circadian rhythm.

The strain homozygous for the inversion (2R*b*/2R*b*) was also independently observed and reported in 1984 in individuals from Karachi[14]. They reported many offspring of a single female that were homozygous for the 2R*b* inversion, which is implicated in resistance to alphamethrin. Alphamethrin was coincidently approved by WHO for adult malaria control around 1981-1985[15]. They observed and reported several inversions across the genome of *An. stephensi* by studying polytene chromosomes from ovarian nurse cells from various strains and locations. In this report, a photomap of both standard and inverted configurations of the chromosomes for all inversions was documented. Using visible loop formation near the inversion region and assigning banding patterns, they were able to identify/validate a number of inversions including three from the 2R arm, two from the 2L arm, three from the 3R arm and several from the 3L arm along with approximate breakpoints based on cytobands. There was no inversion reported in the X chromosome. They also demonstrated that, except for a few inversions, a significant number of inversions cataloged in the wild population, were lost during the first few generations of rearing in the lab. However, among those that were persistent for many generations of rearing in the laboratory are 2R*b*, 2L*c,* 3R*b* and 3L*b*.

More recently, Kamali *et al*.[4] used 12 microsatellite markers in conjunction with a collection of previously-known breakpoints to correlate inversion status with phenotypes. In this report, using microsatellite markers on polytene chromosomes, it was shown that the E7T microsatellite marker, which maps near the 14C band within the 2R*b* inversion, is diverse in the three forms of *An. stephensi* including type, intermediate and mysorensis.

The most extensively-studied malaria vector to date is *An. gambiae*. A number of very well-characterized inversions in *An. gambiae* and their association with important traits, highlights the importance of studying inversions in malaria vectors at molecular levels[11–12].

The characterization of major inversions in *An. gambiae*, such as 2R*b* and 2L*a*, offer a rich source to establish genotype-phenotype association, by synteny, in other malaria vectors, such as *An. stephensi*. The two breakpoints of the 2R*b* inversion are reported to contain repetitive regions, making it prone to structural and sequence level instability[16]. In the case of the 2L*a* inversion, by revealing the structure at the breakpoint it is proposed that this inversion has descended from a single event[17]. Furthermore, the 1000 *An. gambiae* genome project also enabled extraction of SNP signatures useful in the association of 2R*b* inversion genotypes to the respective phenotypes[18]. Cost effective technologies, such as RFLP-PCR, are being explored to type 2R*b* signature SNPs in *An. gambiae* from a large number of individuals[19].

Here, we report the genome assembly of a type-form of *An. stephensi* collected from a construction site in Annanagar, Chennai, India, (IndCh), which was reared in the laboratory since 2016. Unlike the genomes of the two other strains reported for *An. stephensi*, the IndCh strain is homozygous for the standard form of the 2R*b* chromosome configuration (2R+^b^/2R+^b^), allowing us to extract segregating signatures required for the identification of candidate genes responsible for the expansion range of this important vector. Furthermore, the assembly of IndCh reveals the extent of genomic changes in *An. stephensi* during expansion of commercial activity in this subcontinent since the UCI and STE2 strains have been collected.

## RESULTS

### Assembly of the genome of IndCh strain

Using DNA extracted from 60 individual iso-females homogenized by sib mating over five generations, we generated 60X coverage of PacBio reads using Sequel 1 technologies for an average length of 9000 bp and 100X coverage of Illumina reads. PacBio reads were assembled using multiple tools, including CANU[20] and FALCON[21] to get contig-level assemblies with L50 values of 20 and 28, respectively. The two assemblies were merged using Quickmerge[21] to obtain an improved contig-level assembly with a L50 of 8. The merged assembly was corrected for sequencing errors using two rounds of Arrow and Pilon polishing[22] each. The statistics of these assemblies are listed in Table 1.

**Table 1:** Statistics of the genome assembly of the IndCh An. stephensi strain

| **Assembly Stage** | **No. of contigs** | **Longest Contig (Mbp)** | **N50 (Mbp)** | **L50** | **Assembly Size (Mbp)** | **Average Contig Length (Kbp)** | **Median Contig Length (Kbp)** |
| --- | --- | --- | --- | --- | --- | --- | --- |
| **Canu Assembly** | 1,683 | 10.7 | 3.0 | 20 | 219.8 | 130.6 | 10.9 |
| **FALCON Assembly** | 1,250 | 9.7 | 1.8 | 28 | 231.1 | 184.9 | 21.2 |
| **Merged and Polished** | 1,010 | 26.5 | 8.5 | 8 | 225.8 | 223.6 | 19.0 |
| **HiC Scaffolds** | 962 | 35.9 | 21.4 | 4 | 226.9 | 236.0 | 18.6 |

In order to select the most relevant publicly available HiC dataset to build scaffolds for the IndCh strain, an attempt was made to establish the 2R*b* inversion status for the IndCh assembly. For this, sequences of physical markers corresponding to chromosome 2R arm from Jiang *et al.*[6] were aligned against the merged, contig-level assembly mentioned above. The markers spanning the breakpoints for the 2R*b* inversion were distributed on 2 different contigs (contig-982 and contig-2). The linearity of the markers 11A to 12B (left breakpoint) and 16C to 17A (right breakpoint) were both found within the two contigs shown in Fig. 1C, indicating that IndCh supports the standard configuration of 2R*b*. There were no contigs supporting the inverted form of 2R*b* in which one expects 11A followed by 16C on one side and 12B followed by 17A on the other. Furthermore, observing polytene chromosomes from ovarian nurse cells suggests lack of heterozygosity in the 2R arm (Fig. 1B) in 96% of the sixty-one insects bred over 35 generations after the iso-females were selected for sequencing. Haplotype phasing of the assembly of the IndCh strain was done using a set of primary contigs and a set of associated contigs from FALCON and FALCON-unzip[23] tools, respectively. The lack of a double peak in the IndCh phased contigs (Fig. 1A) along with other data mentioned above, supports that IndCh has the standard homozygous form of 2R*b*.

**Fig. 1:**
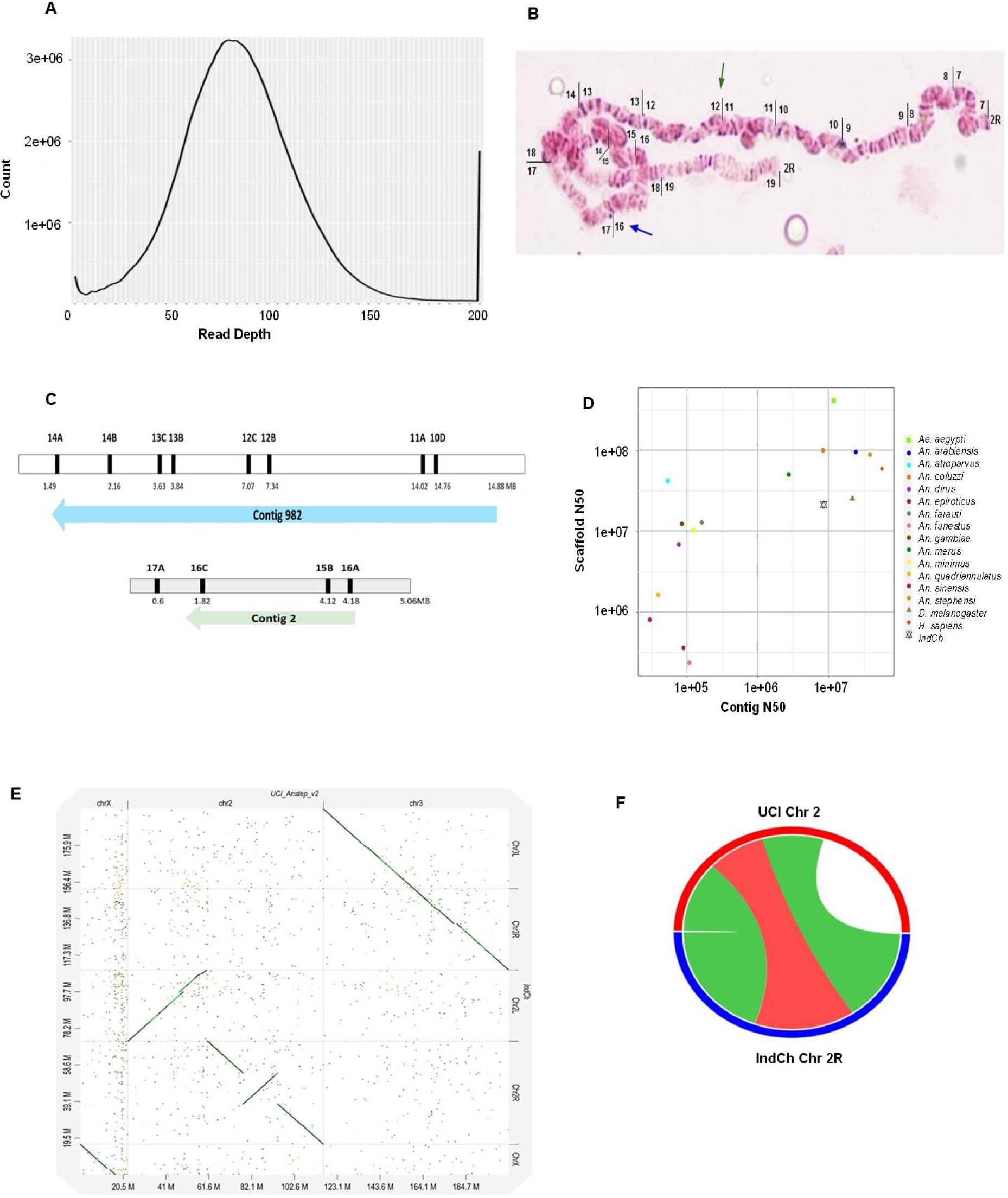
(A) Coverage analysis for the assembly of the IndCh strain with single hump instead of double expected for heterozygosity in the 2Rb inversion. (B) Photomap of polytene chromosomes from the IndCh strain showing the standard configuration of 2Rb in 96% of individual mosquitoes. The green and blue arrows point to the 2Rb breakpoints of the standard form. (C) Two contigs in support of the standard form of 2Rb. Contig-982 (top) supports the left breakpoint and contig-2 (bottom) supports the right breakpoint. (D) Comparison of the assembly statistics with other Anopheles, fly and human genomes. (E) Dot-plot of IndCh assembly against the UCI genome. (F) Synteny between UCI chromosome 2 and IndCh chromosome 2R, with the red part indicating the 2Rb inversion region, the green part indicating the spanning uninverted regions, while the white gap is from the 2L arm of the UCI chromosome.

The HiC reads generated from the STE2 strain were selected for scaffolding the merged contigs of IndCh because the STE2 strain, which is heterozygous for the 2R*b* inversion, contains reads representing the standard form of 2R*b*[3]. A scaffold-level assembly with a L50 of 4 was obtained for IndCh (Table 1). Scaffolds were assigned both the positions and orientations on the chromosomes using physical markers[6] and were stitched into pseudo-chromosomes (Supplementary Fig. 1). Fig. 1D compares the quality and completeness of the assembly reported here to those of the reported genomes of other *Anopheles* species, the fly and the human, suggesting that the IndCh assembly is comparable to the other high quality genomes on both the contig and scaffold levels. A pairwise synteny between the IndCh and UCI assemblies clearly shows the 2R*b* inversion in the UCI genome, when compared to the reported assembly of IndCh (Fig. 1E). Fig. 1F shows synteny between the 2R arms of the IndCh assembly and the genome of the UCI strain.

### Interrogation of 2R*b* breakpoints

With high-resolution genome assemblies from two different homozygous configurations of the 2R*b* and availability of HiC data from two strains with different 2R*b* configurations, an attempt was made to define the 2R*b* breakpoints for *An. stephensi* at the sequence-level for the first time. The rationale was to map the HiC reads from a given strain onto the assembly with the opposite 2R*b* configuration to produce off-diagonal butterfly structures, the center of which will represent the breakpoints on the respective assembly, as shown in Fig. 2. For example, HiC data from the STE2 strain, which is heterozygous for the 2R*b* inversion, but contains reads representing the standard 2R*b*, was mapped to the genome of UCI strain, which is homozygous for the 2R*b* inversion (Fig. 2A). In this case, the read-pairs making the butterfly support the standard form of 2R*b*. On the other hand, HiC data from UCI that contains reads supporting only the homozygous inverted form (2R*b*/2R*b*) was mapped onto the assembly of IndCh, which is homozygous for the standard form (2R+^b^/2R+^b^), creating two very sharp butterfly structures corresponding to two breakpoints in the IndCh assembly (Fig. 2B). The coordinates of the center of the butterfly in both contact maps were deciphered to a maximum resolution of 5 Kbp per dot by enlarging the contact map. Using this approach, the loci of 2R*b* breakpoints in the UCI genome were estimated to be between chr2:55265000-55270000 on the left, and chr2:71795000-71800000 on the right (Fig. 2A). Likewise, the loci of the 2R*b* breakpoints in the IndCh assembly were estimated to be between chr2R:21510000-21515000 on the left, and chr2R:38085000-38090000 on the right (Fig. 2B).

**Fig. 2:**
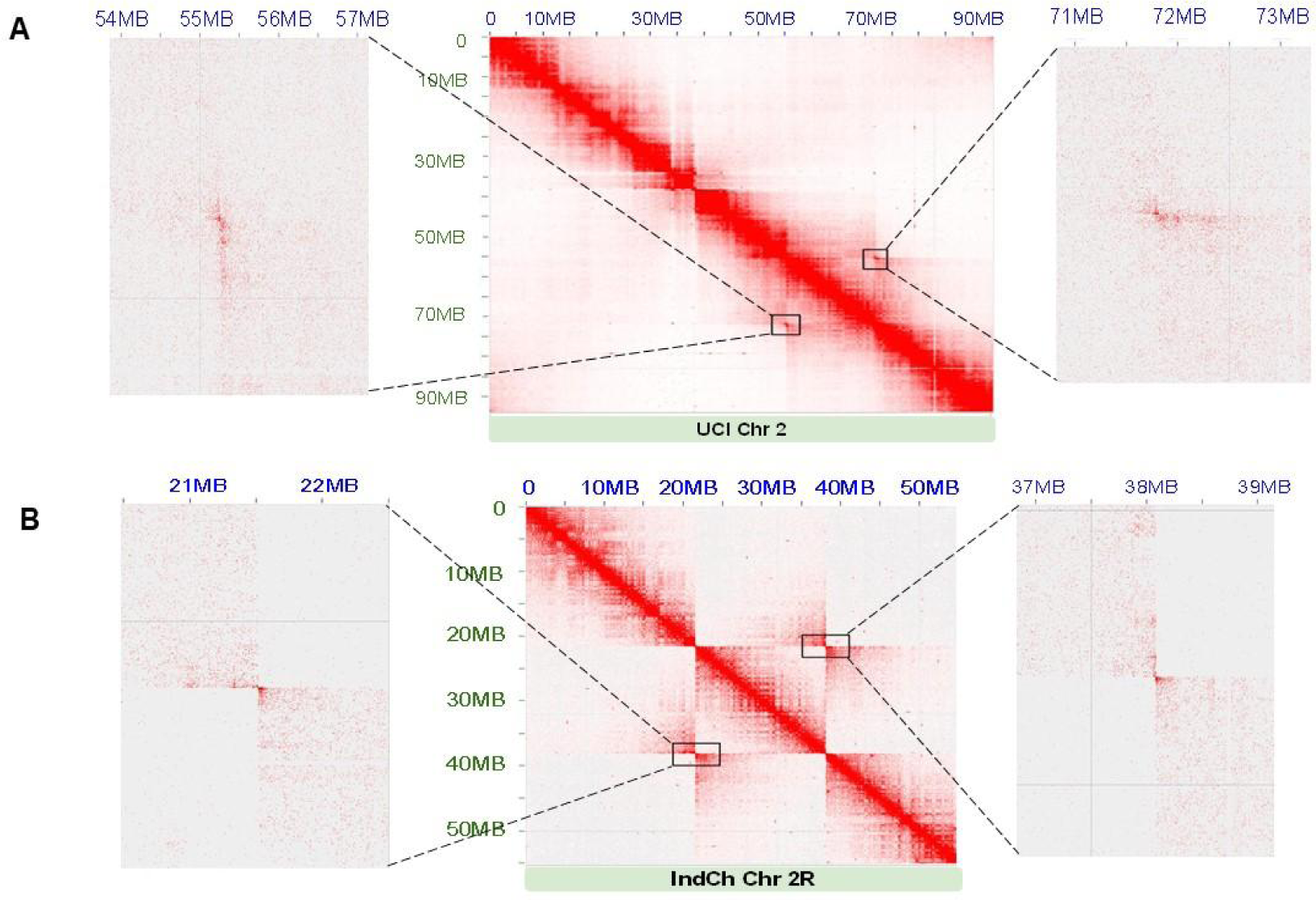
(A) Left and right breakpoints obtained by mapping STE2 HiC against the chromosome 2 of UCI. The black box represents the left and right breakpoints located between the ranges of 55,265,000-55,270,000 bp and 71,795,000-71,800,000 bp, respectively, in the UCI genome. Adjacent to the contact map are the enlargements of the butterfly structures, for a resolution of 5 Kbp per dot. (B) Left and right breakpoints obtained by mapping the UCI HiC against the chromosome 2R of IndCh. The black box represents the left and right breakpoints located between 21,510,000 to 21,515,000 and 38,085,000 to 38,090,000 bp, respectively, in the IndCh assembly. Adjacent to the contact map are the enlargements of the butterfly structures for a resolution of 5 Kbp per dot, respectively, in the IndCh assembly.

In order to further refine the loci of the breakpoints, the error-corrected PacBio reads of IndCh were mapped onto the chromosome 2 of the UCI genome. Since the UCI and IndCh strains are of opposite genotypes, one would expect a gap among IndCh reads near the UCI breakpoints, (Supplementary Figs. 2A and 2B, top panel). Based on the location of the gap, the locus of the left breakpoint was refined to chr2:55256667-55257873, which is 10 Kbp or two dots (5 Kbp each) left of the estimated breakpoint from the contact map. Similarly, based on the location of the gap, the right breakpoint was refined to chr2:71807500-71808000, which is also 10 Kbp or two dots right of the estimated breakpoint locus. Furthermore, mapping of Illumina reads from IndCh onto the genome of UCI, confirmed the refined breakpoint loci (Supplementary Figs. 2A and 2B, bottom panel). On the other hand, in support of the authenticity of IndCh assembly near the breakpoints, both error-corrected PacBio and Illumina reads from IndCh showed no gaps (Supplementary Figs. 2C and 2D), thus validating the assembly near the standard 2R*b* breakpoints.

### Characterizing breakpoints

The IndCh assembly was mapped to the genome of UCI using ‘unimap’ (https://github.com/lh3/unimap). There are three blocks from the 2R arm of UCI genome with extensive homology with IndCh assembly, as shown in the schematic Figs. 3B and 3D. On the two sides of the 2R*b* inversion, the region ∼71.8 to 93.7 Mbp of UCI 2R arm maps to the region from the beginning of the 2R arm of IndCh through 21.5 Mbp. Similarly, the region from ∼38.3 to 55.3 Mbp of the UCI 2R arm maps to a region from ∼38.1 to 55.0 Mbp of the IndCh assembly. Within the 2R*b* inversion, near the locus ∼66 Mbp in UCI and ∼32 Mbp in IndCh, there are significant duplication and minor inversions in IndCh, as shown in Fig. 3A. The homologous parts of the 2R*b* inversion in both the assemblies are around 16.5 Mbp and on the two sides of the inversion, the homologous 2R arm is nearly 17 Mbp and 21 Mbp in both assemblies. On the other hand, near the breakpoints, thousands of extra bases are found in the genome of UCI strain with very short stretches of corresponding sequences in the IndCh assembly, as shown between blocks in Fig. 3D. Interestingly, these extra bases are filled with repeat elements, as shown in Fig. 3E.

**Fig. 3:**
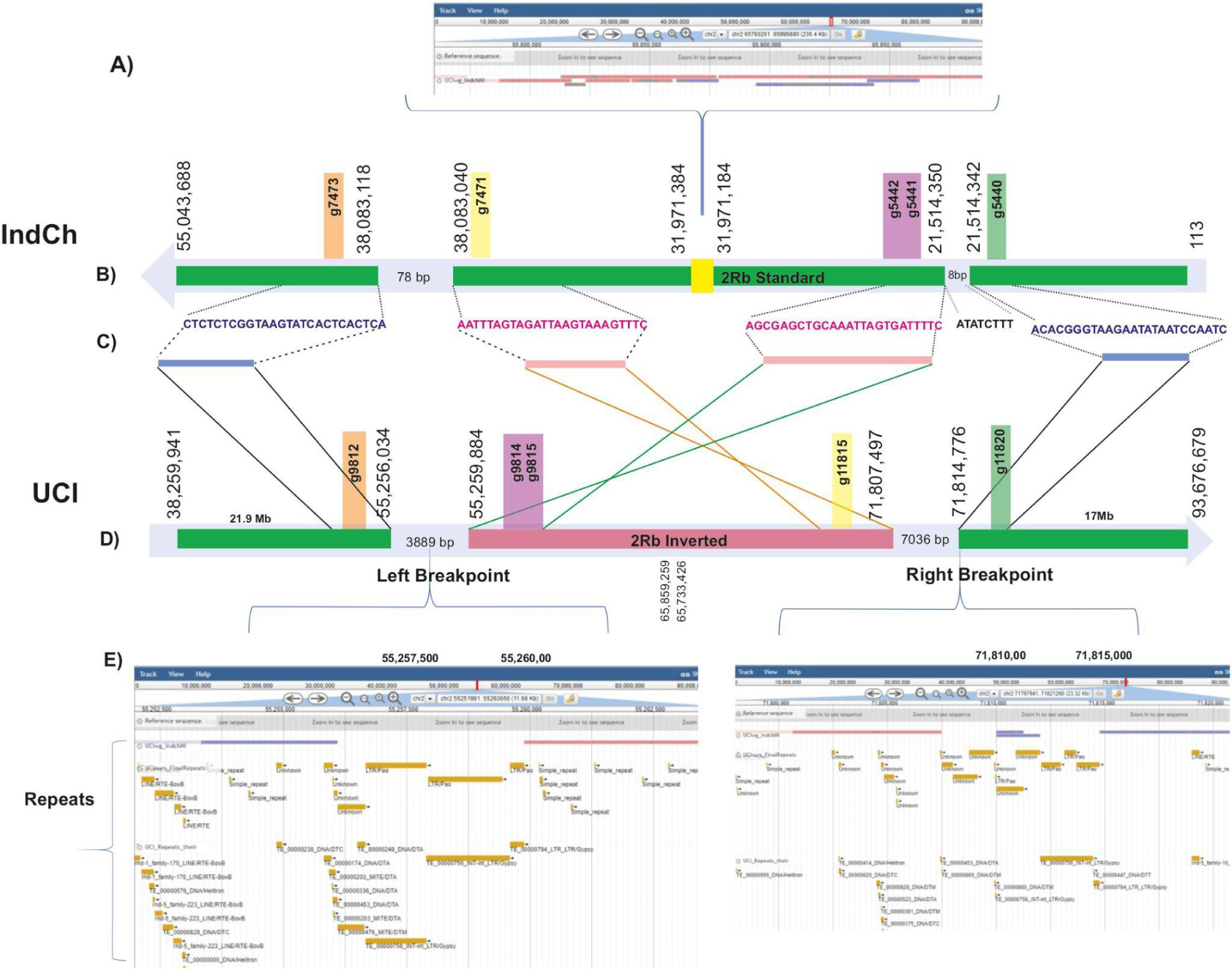
(A) A schematic of unimap of the IndCh assembly on the UCI genome showing the overlapping segments in the middle of the 2Rb inversion. (B and D) IndCh and UCI blocks, respectively, that are homologous with chromosome coordinates at the start and end of each block. (C) Highlights the sequences near the breakpoints shown by blue boxes (standard orientation) and the pink boxes (inverted orientation) refer to the two breakpoints/gaps between the three homologous blocks shown on the browser, (E) Repeat elements in the gap between breakpoints in the UCI assembly.

Fig. 3C shows alignment of the homologous sequences near the breakpoint from IndCh to the corresponding region in the genome of UCI determined using the BLASTN tool. We also find orthologous genes (color coded in Figs. 3B and 3D) on both sides of the two breakpoints that are consistent with the inversion phenotype. Fig. 3C highlights the sequence near the breakpoints. The sequences spanning the breakpoints in the IndCh assembly are given in the Supplementary Text 1 and 2 and have been validated using PCR. As shown in the Supplementary Fig. 3A, one of the primer pairs amplified the left breakpoint from individuals from outgrown IndCh population by 35 generations and another primer pair amplified the right breakpoint (Supplementary Fig. 3B) only from the parent population from which the IndCh iso-female line was originally generated (Fig. 8).

### Genes within the 2R*b* locus

The 2R*b* locus spans 16.56 Mbp and has 1,353 predicted genes. Interestingly, these genes include ACE1, tandem clusters of GST and CYP450 paralogs implicated in insecticide resistance. This locus also includes 3 paralogs of GILT genes, one of which, GILT3, is expressed in salivary gland and slows the mobility of *Plasmodium* after it is injected into the bloodstream of the human host. The GILT3 gene in IndCh is structurally disrupted with respect to the UCI strain due to a 20 bp insertion. It is likely that this disruption is responsible for the higher vectorial capacity of the IndCh strain.

We looked into SNVs, indels and structural variants (SVs) within genes in IndCh, when compared to the UCI strain. A total of 1.6 million SNVs are found in chromosome 2 of IndCh compared to the UCI genome, of which 19,541 create missense mutations and 3,599 fall within the 2R*b* locus. Interestingly, we found 4 missense mutations within the ACE1 protein alone, of which R480L is most likely to impact its function (Fig. 4E). One of the Cyp6a2 transcripts also has a non-consevative, amino acid substitution, A61E, in the IndCh assembly (Fig. 4D). Using the publicly-available developmental transcriptome data, we also showed the expression patterns of genes within the 2R*b* locus that are implicated in insecticide resistance, such as GST and Cyp450 gene clusters. Some of these genes are upregulated in the larva stage in STE2 strain, suggesting their role in insecticide resistance (Fig. 4A, 4B).

**Fig. 4.**
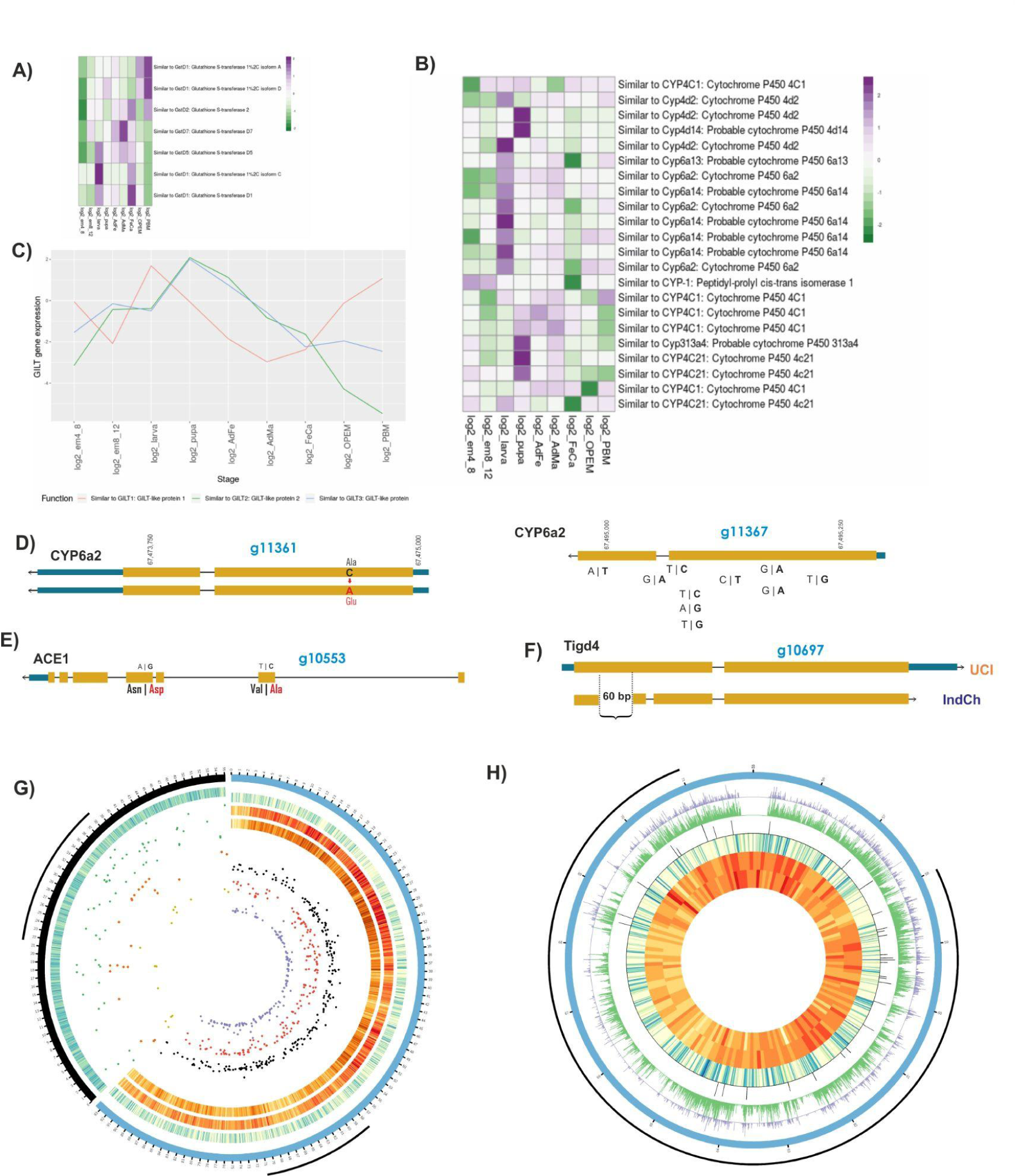
top: (A) Expression profiles of GST paralogs from within the 2Rb locus across developmental samples. (B) Expression profiles of Cyp450 paralogs from within the 2Rb locus across developmental samples. (C) Expression profiles of GILT paralogs from within the 2Rb locus across developmental samples. (D) Missense SNP in Cyp6a within the 2Rb locus. (E) Two of the four missense SNPs in the ACE1 gene within the 2Rb locus. (F) The gene Tigd4 shows a 60 bp deletion in the middle of the first exon in the IndCh assembly. (G) The outer black arc represents IndCh Chr2R and the blue arc represents UCI Chr 2. (From inside to outside, from left to right) - The heat map represents the gene density, green dots are insertions and the orange dots are the Gypsy elements intersecting with these insertions in the IndCh strain. The red dots are the LTR-Gypsy elements in the UCI strain and the black dots are the deleted SVs. The heat maps show the density of deletions, insertions and genes respectively. (H) (From outside to inside) The black arc represents the 2Rb region in the UCI strain. The blue circle is chromosome 2 of the UCI strain from 55 to 72 Mbp. The purple peaks represent 3,781 significant SNPs with a p-value of 0.0005 among the candidate SNPs for 2Rb, the green peaks represent the 22,650 candidate SNPs for 2Rb, while the black peaks are the densely segregating SNPs (277) from the obtained blocks (7 or more SNPs in a 1,500 bp region). Followed by these are the tracks for the density of genes, insertions and deletions, respectively.

There are 3,881 structural variants (SVs) within the 2R*b* inversion locus in the IndCh assembly compared to the UCI genome, including 1,905 deletions, 1,727 insertions, 21 CNVs, 185 breakends, 6 inversions, 29 insertion duplications and 8 split duplications. The two coloured heat maps (Fig. 4G) shows the number of deletions and insertions in IndCh compared to chromosome 2 of UCI (outer blue circle in Fig. 4G), suggesting that the density of SVs within the 2R arm is lesser than that in the 2L arm of UCI. Deletions in IndCh with respect to the UCI genome alone account for 3.8% of the genome (Supplementary Fig. 4). In order to visualize the impact of structural variants, it was necessary to liftover the predicted IndCh genes to the UCI genome assembly. The diversity in the two strains posed a major challenge with regards to the liftover. This was done using a home-grown tool, which allowed liftover of 80% of the genes, an example of which is shown in Supplementary Fig. 5.

There are 30 SVs that intersect with exons and have the potential to alter functions of 18 genes within the 2R*b* locus, including some genes of unknown function. The functions of the remaining 14 genes were checked along with the presence of more copies, or their paralogs, throughout the genome. Among a few with interesting structural changes are Lin37, MCM3AP, Tigd4, Nop58 and Tango2 genes. The 5’UTR of Lin37 appears to be translocated from the 5’ UTR of MCM3AP in the UCI genome. This region has no repeat element, but is as long as 5 Kbp in size. So this translocation has the potential to alter the structure, function or expression of both Lin37 within the 2R*b* loci and the MCM3AP gene in chromosome 3. The MCM3AP is a gene involved in the transport of mRNA to the cytoplasm. It acetylates MCM3, which then takes part in DNA replication[24–26]. Lin37 gene is involved in cell-cycle checkpoints and is part of the DREAM complex activated whenever the G1-S phase, cell-cycle checkpoint protein, retinoblastoma, loses function.

Among other structural changes are the first exon of the gene, Tigd4, with a 60 bp deletion in the middle of the first exon in the IndCh assembly. The Nop58 gene[27] in IndCh shows one less exon as compared to that in the UCI genome because of a premature stop codon. Since both Tigd4 and Nop58 have structurally uninterrupted paralogs, CENPB and Nop56, respectively, any negative impact of these deletion/termination on the function may, perhaps, be compensatory. The Tango2 gene in IndCh is a copy number variant with one copy resulting in a frameshift variant with loss of the start codon. A depletion in the protein product of this gene has been shown to cause fusion of Golgi bodies and endoplasmic reticulum (ER) in *Drosophila*[28]. Should the Golgi and ER fuse, there will be a metabolic crisis in the cells of the organism. However, in the IndCh assembly, the duplicate copy of the entire Tango2 gene is intact, suggesting active evolution of this gene in IndCh.

The predicted structural variants in the IndCh assembly also include large deletions and insertions spanning more than 5 Kbp within the body of the genes, which match with LTR-Gypsy elements in the genome of UCI (Supplementary Table 1). For example, there are 244 deletions larger than 5 Kbp in chromosome 2 of IndCh (black dots against UCI chromosome 2 in Fig. 4G), of which 139 match with the LTR-Gypsy elements in the UCI genome (shown by red dots against the chromosome 2 of UCI in Fig. 4G). In order to show the number of LTR-Gypsy inserted in IndCh, Fig. 4G is complemented with the 2R arm of IndCh, shown as the black outer ring. The insertions that are greater than 5 Kbp in the IndCh strain are marked with green dots and the intersecting LTR-Gypsy elements are shown in orange dots. For example, these LTR-Gypsy elements were also observed to be deleted from the IndCh assembly and IndCh PacBio reads when compared with the UCI genome (Supplementary Fig. 6).

### The 2R*b* inversion signature

The reported genome assemblies of the homozygous, standard form of the 2R*b* (2R+^b^/2R+^b^) along with the genomes of the other strains, STE2 and UCI, with heterozygous (2R+^b^/2Rb) and homozygous (2R*b*/2R*b*) forms of the inversion respectively, were used to identify a unique, 2R*b*-specific SNP signature for typing purposes. As shown in Fig. 4H (concentric 5th circle from inside->out in green), 22,650 SNP positions from the 2R*b* locus with homozygous alternate alleles in the IndCh assembly, with the corresponding heterozygous alleles in the STE2 genome and homozygous-reference alleles in UCI genome were selected (schematic Fig. 7). This set of SNPs was used as a signature to perform unsupervised clustering of 15X coverage of WGS data from 62 individuals from four laboratory reared populations and 20 individuals from two wild populations, to produce the cluster shown in Fig. 5A. By clustering close to the IndCh strain, ∼65% of individuals from each population show the homozygous standard form, and the remaining 35% show the heterozygous configuration for the 2R*b* inversion by clustering with the STE2 strain, which is known to be heterozygous. Fig. 5B shows a phylogenetic tree assigning 2R*b* genotype to other strains for which draft genomes are available. Accordingly, the strain SDA500[29], is heterozygous for the 2R*b* genotype and IndInt strain, an intermediate form of *An. stephensi* for which a genome is currently being assembled in-house (unpublished data), is homozygous for the standard form, similar to IndCh strain. As a negative control, a similar strategy was used to extract SNPs from the 2L arm of similar length, which clusters all 82 samples into a single cluster, albeit close to the IndCh genome (Fig. 5E). This would suggest that, except for the 2R*b* inversion locus, all 4 populations are much closer to the IndCh genome with respect to fixed SNPs in the 2L arm.

**Fig. 5:**
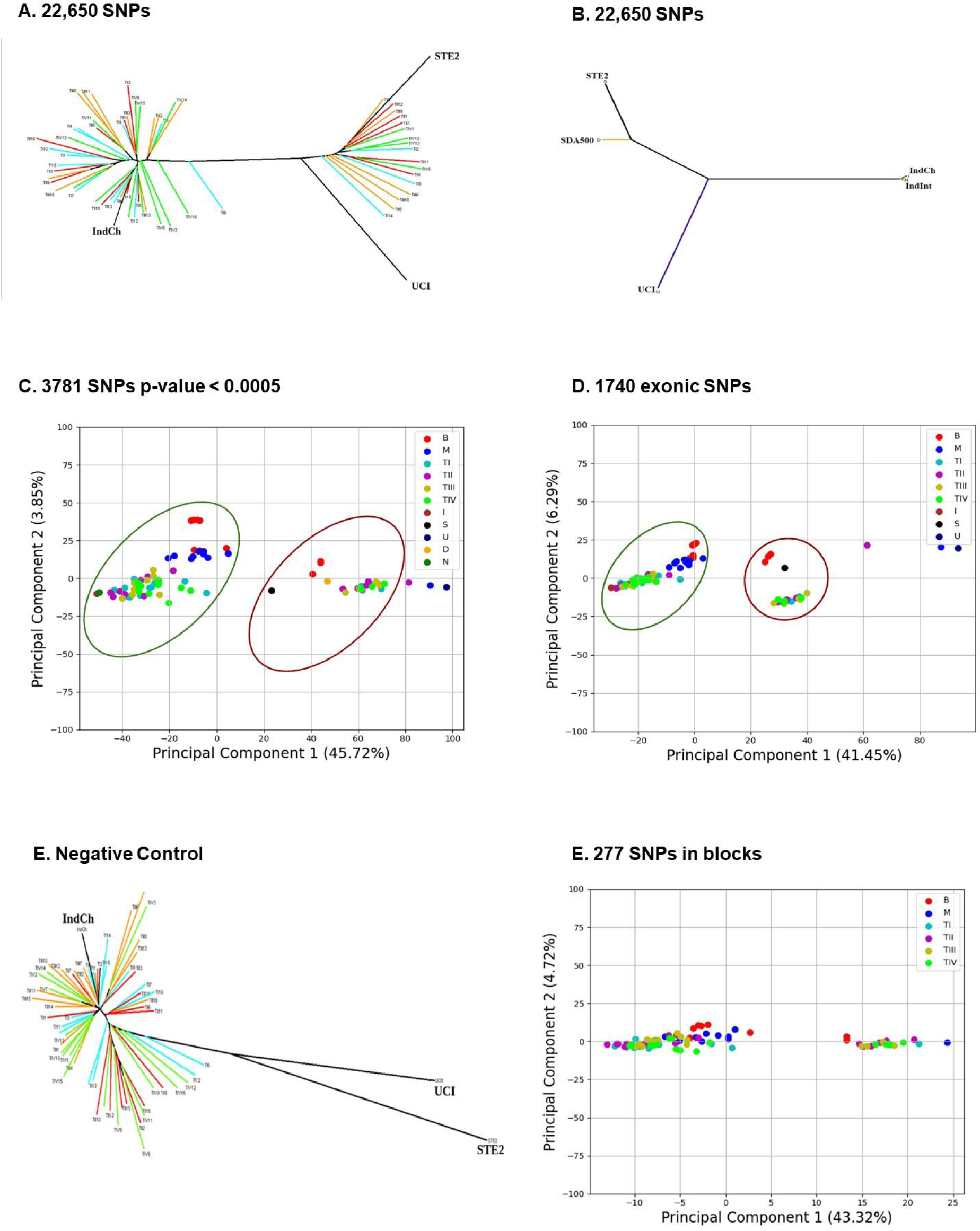
(A) Phylogenetic tree with 22,650 SNPs for 62 laboratory samples along with the three reference genomes. (B) Phylogenetic tree with 22,650 SNPs for five genomes. (C) PCA of 62 lab samples using 3,781 high-significance SNPs obtained after supervised clustering with 22,650 SNP and filtering SNPs with p-value of <0.0005 for laboratory samples with the inclusion of the IndCh, STE2, UCI, SDA500, IndInt genomes. (D) PCA plot of 1,740 candidate SNPs found in the exonic regions of the genome. (E) Negative control using 17,106 candidate SNPS from chromosome 2L of similar lengths. (F) PCA using 277 SNPs from 31 blocks among the candidate SNPs including wild samples.

The 22,650 SNPs were assigned cluster numbers based on the initial unsupervised clustering from Fig. 5A to perform supervised clustering to extract 3,781 SNPs that were most significant with p-values of <0.0005 (Fig. 4H, concentric 6th circle from inside->out in blue). The PCA of the 82 samples, using the 3,781 SNPs, show two distinct clusters revealing segregation of individuals by 2R*b* genotype (Fig. 5C) in line with the cluster assignment in Fig. 5A. Out of the 3,781 highly-significant SNPs, 277 SNPs from 31 SNP-blocks containing clusters of more than 6 SNPs within stretches of 1500 bases, were identified (Fig. 4G, concentric 4th circle from inside->out in black). The 277 SNPs found in WGS data continue to cluster the 82 samples, including lab and wild into two clades (Fig. 5F). In order to assign 2R*b* genotypes for individuals directly from transcriptome data, these 1740 SNPs out of the 22,650 SNPs from within exons, also segregate the 82 individuals into 2R*b* genotypes (Fig. 5D). It should be mentioned that 83% synonymous and 77% of missense mutations from the 1740 exonic SNPs are present in publicly available RNA-Seq data across developmental stages for the STE2 strain, allowing assignment of the 2R*b* genotype directly from transcriptome sequencing data to connect the impact of 2R*b* genotype on gene expression.

## DISCUSSION

Here, we report the assembly of the genome of a novel strain of *An. stephensi* (IndCh), which was collected in 2016 from a construction site in Annanagar, Chennai, India and has since been reared in the laboratory. The genome assembly of this strain completes the trilogy of configurations with respect to a most frequent inversion, reported since 1984, spanning 16 Mbp with more than 1300 genes in the middle of the 2R arm of *An. stephensi* chromosome, implicated in adaptation to insecticides, behaviour and climate. The complementarity of the IndCh genome with that of the UCI strain reported recently, allowed virtual reconstruction of a heterozygous 2R arm (Fig. 6). We hypothesize that the thousands of bases inserted at the 2R*b* breakpoints in the inverted form in UCI strain may be required to sterically accommodate loop formation in heterozygous configuration, without disrupting the structures of homologous segments. The right breakpoint with more than 7 Kbp insertion may be more fuzzy leading to repetitive breaks as revealed by the photomap of heterozygous individuals from iso-female lines after 40 generations (data not shown). This is also confirmed by the failure of PCR amplification of the right breakpoint from IndCh from the outgrown iso-female line (Supplementary Fig. 3B). Since the right breakpoint of 2R*b* overlaps with the left breakpoint of 2R*i* locus[30], we conclude that the right breakpoint of 2R*b* may be involved in more than one inversion type and perhaps is in active evolution in this population. However, the right breakpoint could be amplified from the Chennai (TII) laboratory strain, the parent population from which the IndCh strain is originally derived (see PCR in Supplementary Fig. 3).

**Fig. 6:**
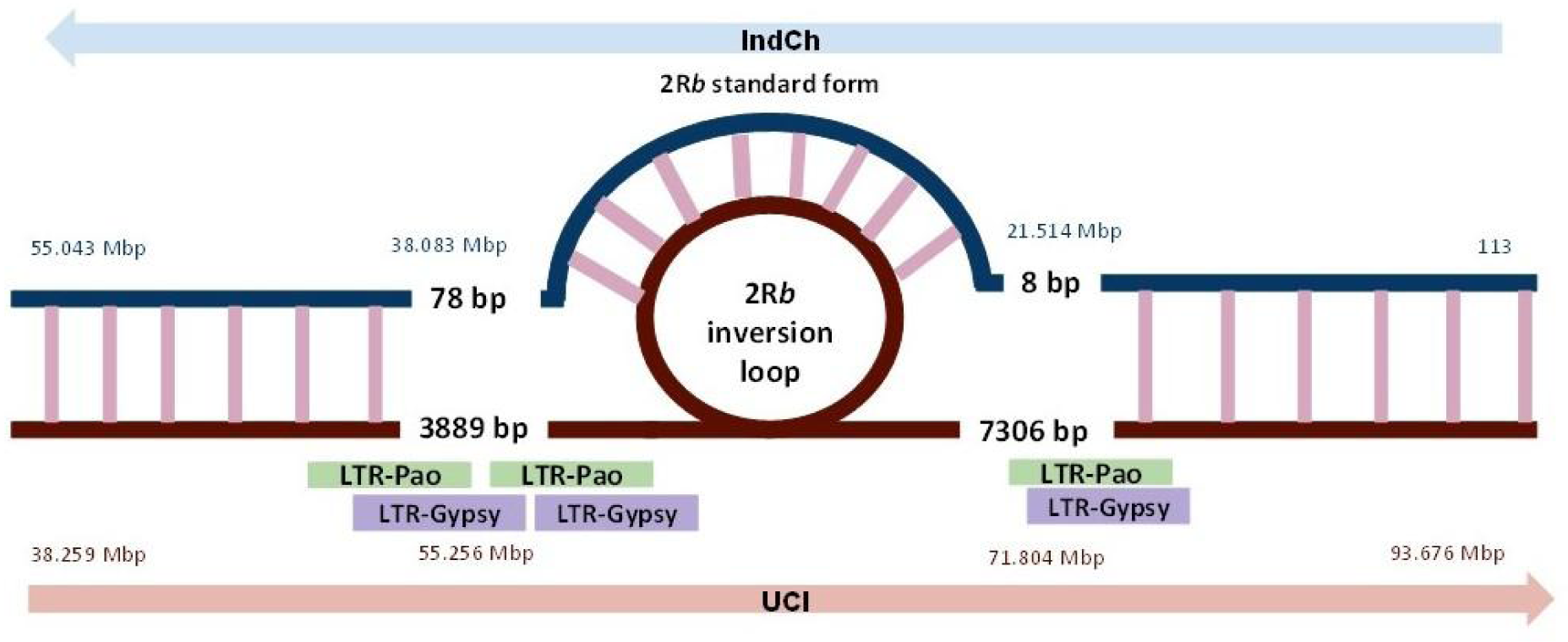
Cartoon of a simulated heterozygous loop formation of the 2R arm of chromosome 2 built using the genome of homozygous 2R+^b^/2R+^b^ form from IndCh (blue) and 2Rb/2Rb form from UCI (maroon). The homologous segments are shown with pink bridges.

**Fig. 7:**
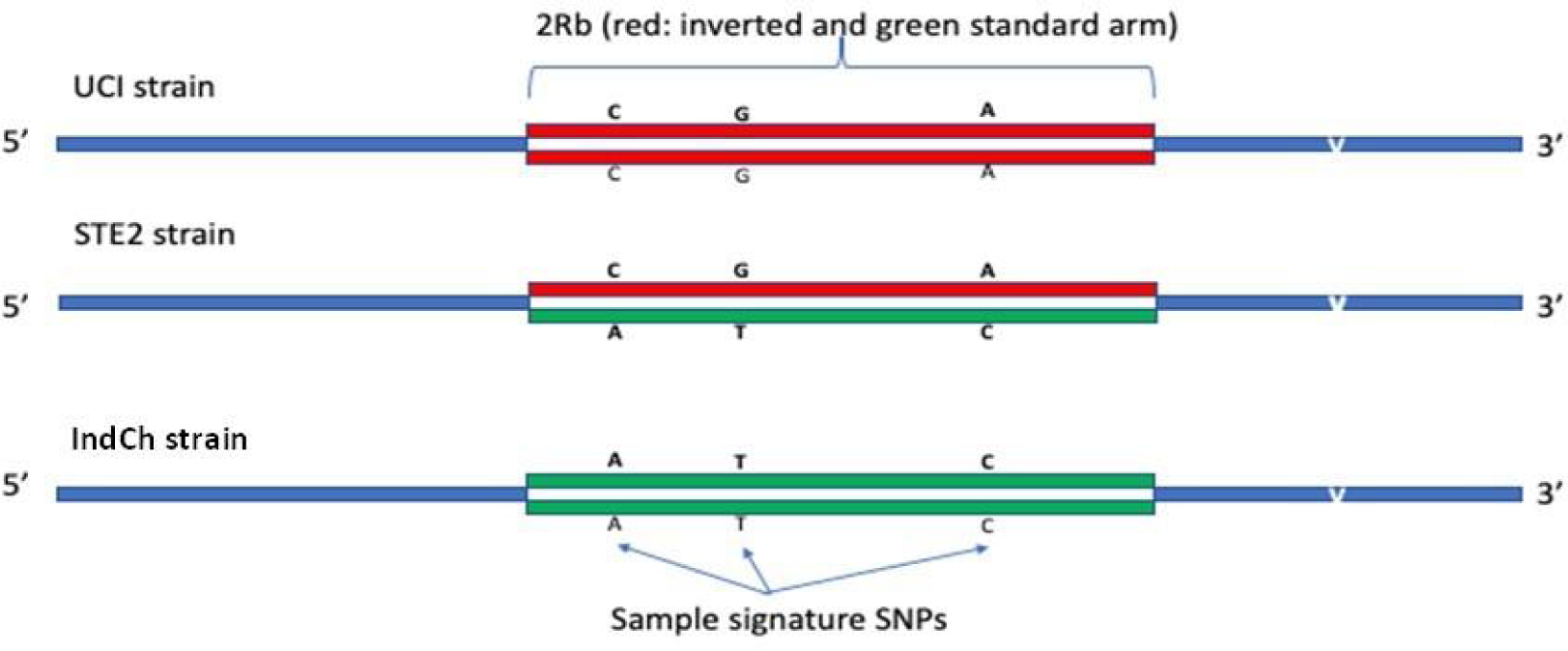
Strategy for selection of SNPs and indels specific for 2Rb standard and inverted forms.

In *An. gambiae,* it was reported that transposable elements (TEs), including the LTR-Gypsy elements, accumulate near the 2L*a* inversion breakpoints[31]. These authors mention that both 50 Kbp regions immediately surrounding each breakpoint in 2L*a* contain a highly significant increase in TE density. In this context, the thousands of bases inserted near the breakpoints in the UCI genome of *An. stephensi*, filled with repeats, suggest that inversions are not just relative arrangement of physical markers, but have bearing in mosquito evolution.

Structural variants caused by repeat elements have been shown to be prevalent among variations of complex traits in *Diptera*[32]. Since retrotransposons, including LTR-Gypsy, impact the expression of genes, insertion and deletion of these elements have been postulated to affect the efficiency of the genes inserted via CRISPR-Cas9 construct in gene-drive technologies[33]. There are a total of 43 predicted genes within the 2R*b* locus in IndCh with LTR-gypsy element inserted/deleted, perhaps, regulating their expression differentially in other genotypes of 2R*b*.

The unusually large number of segregating SNPs identified allows the opportunity to create several subsets of signatures for typing using diverse cost-effective technologies, such as low-density microarrays, ampli-seq, RNA-seq or PCR. The typing of 82 individuals using WGS demonstrate that 30% of individuals within lab-reared and wild individuals maintain heterozygous form of the 2R*b* genotype consistent with the observation reported by Mahmood and Sakai[14] by studying loop formation in polytene chromosomes across generations back in 1984. It should be mentioned that even in individuals from the Chennai population, the source of IndCh strain, 30% display heterozygous forms of 2R*b* (Fig. 5A). Even the outgrown iso-female line used for assembling IndCh, maintained separately over 35 more generations (see Fig. 8), start displaying 3.18% heterozygosity for 2R*b* (Supplementary Fig. 7), suggesting some remnant heterozygosity from the parent Chennai lab population continues to persist.

**Fig. 8:**
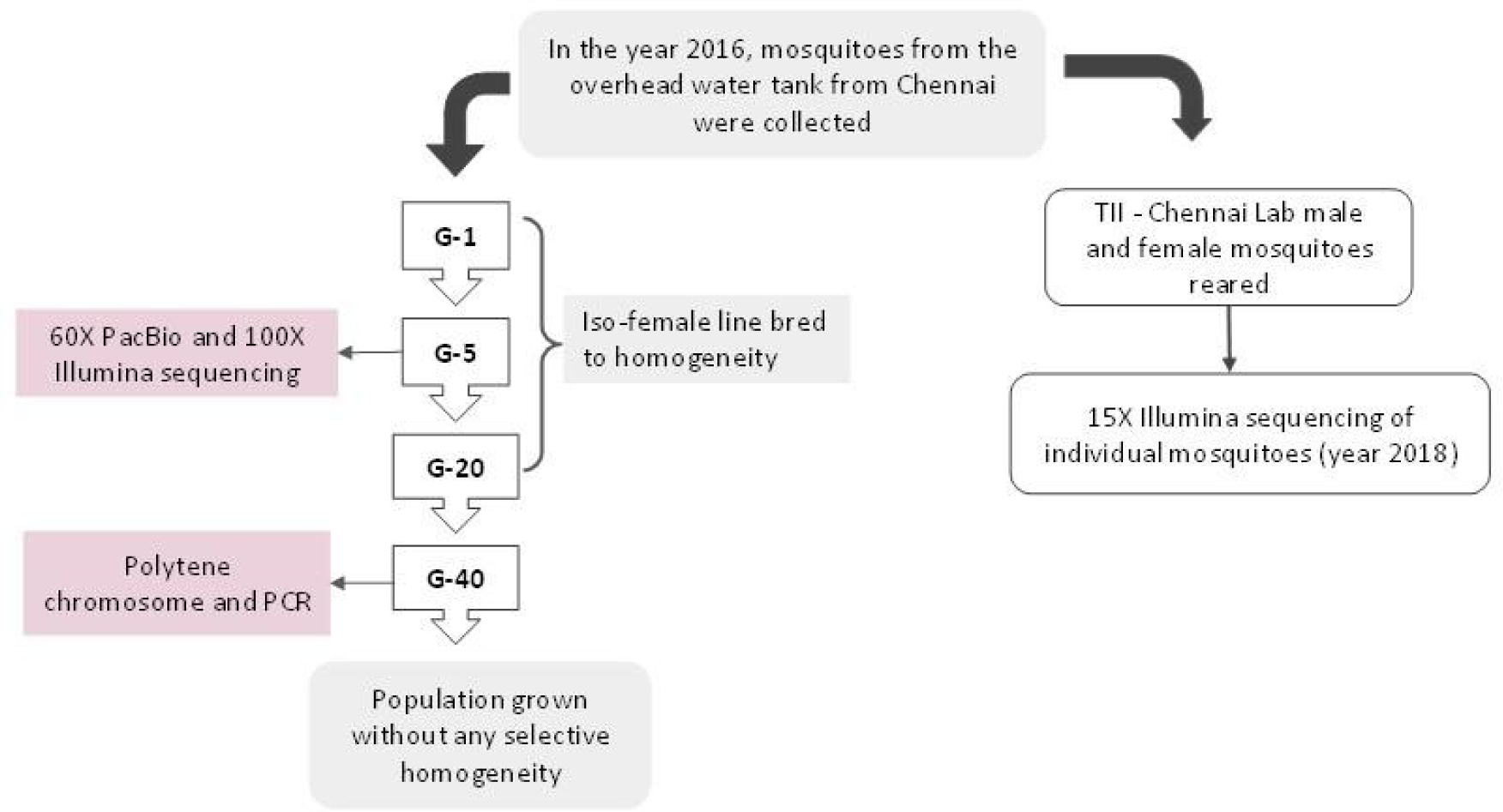
Flow-chart showing the rearing and generation-wise data for Chennai mosquitoes used for raw sequence data, validation using PCR, generating photomap using polytene and analysis.

Given the importance of 2R*b* inversion genotype to adapt to environmental heterozygosity, it is desirable to type directly using transcriptome sequencing, which will allow association of gene expression with inversion genotype. In Fig. 5D we show that the set of 1740 out of the 22,650 segregating SNPs from within exons can be typed using RNA-seq, allowing one to directly associate 2R*b* genotype with gene expression. From the publicly available transcriptome data for adult females[6], 1122 out of the 1740 exonic SNPs can be used for typing. Considering that decades of genomic efforts in *An. gambiae*, a malaria vector extensively studied, only 349 segregating SNPs for the 2R*b* inversion is reported[34].

The clustering of 82 individuals from diverse geographical areas for SNPs from a chromosomal locus used as a negative control cluster in the same clade as IndCh (Fig. 5E), and distal from clades representing STE2 and UCI. Furthermore, a comparison of genome-wide variants among the genomes of the three strains of *An. stephensi* reveals that IndCh has ∼2 million variants against UCI strain, which is 3-fold more than the ∼600K seen for STE2 against the UCI genome, Furthermore, in IndCh, there are 30K missense mutations within the 14K predicted genes reported for UCI genome[9] compared to only 10K missense mutations in STE2 compared to UCI. These observations suggest that IndCh has not only diverged from the other two strains but significant cross breeding may have happened in recent times with increased commerce between major cities. This is even more evident from clustering of individuals from TIII (Delhi-lab) with IndCh, and away from STE2, originally collected from Delhi decades ago.

The complete genomes of multiple strains of *An. stephensi* allows for deciphering genes within other inversion loci, by synteny to completed genomes of other extensively studied malaria vectors. For example, although 2R*b* locus of *An. gambiae,* which is also implicated in insecticide resistance[35] showed no synteny with the 2R*b* locus in *An. stephensi* (Supplementary Fig. 8), the 2R*i locus* of *An. stephensi* is partially syntenic to 2R*b* of *An. gambiae*, both of which are implicated in DDT resistance. Also, the synteny between 2L*a* locus in *An. gambiae* with the 3L*i* and 3L*h* loci in *An. stephensi* may lead to the identification of genes responsible for *Plasmodium* susceptibility in *An. stephensi*.

## CONCLUSION

To our knowledge this may be the first instance where all three genotypes of any inversion in any *Anopheles* species have been deciphered at high-resolution. This has allowed identification of large number of segregating SNPs using comparative genomics and, for the first time, one can type 2R*b* status using transcriptome sequencing, a critical step in the identification of candidate genes from the more than one thousand genes within this locus that may be responsible for traits like insecticide resistance, circadian rhythm and adaptation to climate. We believe that the IndCh genome reported here is a more relevant reference for malaria management in urban settings because there is significant divergence in IndCh collected in 2016 from the earlier strains. Furthermore, our data suggests that increased commerce between various cities in India, in recent times, may have caused a convergence in genotype reflected in IndCh that is most fit to urban settings. The genome trilogy with respect to 2Rb genotype reported here and findings from comparative genomics has put *An. stephensi* genome resource, a grossly understudied malaria vector until recently, on par with the genome resources accumulated over two decades for the most extensively studied malaria vector *An. gambiae*.

## METHODS

### Collection and maintenance of mosquito colony

*An. stephensi* was collected from Anna Nagar, Chennai city, India and is being maintained in the insectarium of Tata Institute for Genetics and Society (TIGS, India). For routine maintenance larvae are given food prepared by mixing Brewer’s yeast powder and dog food (Pedigree brand chicken and vegetable mix) at a ratio of 30:70[36]. Pupae are bleached with 1% sodium hypochlorite for 1 min^35^ and kept in a mosquito rearing cage (Bugdorm-4S3030) (W32.5 x D32.5 x H32.5 cm) for eclosion. The emerging adults are fed on a mixture solution of 8% sucrose, 2% glucose mixed with 5% multivitamin syrup (Polybion LC^®^)[36]. Standard membrane feeding system with modification was followed as described in Gunathilaka *et al.*[37]. On 3^rd^ day eggs are collected and allowed to hatch out into larvae in rearing trays (Polylab, Catalogue no. 81702) (L39 x B30 cm). The larval stage of mosquito is completed within 12-14 days. Adults are maintained in the insectarium at 28±1°C temperature and relative humidity 75±5% with a photoperiod of 12:12h dark and light cycles.

Biosafety approval for mosquito maintenance facility (approval Ref. No.TIGS 2nd IBSC Oct 2018) and Institutional ethical approval for use of human blood for mosquito feeding were obtained (approval Ref. No. inStem/IEC-12/002).

### Establishment of iso-female lines

IndCh iso-female line of *An. stephensi* was generated from the above-mentioned mosquito colony following the protocol of Ghosh & Shetty[38]. Blood meal was given to 5-day old adult females. Twenty-five gravid females were selected and kept separately. On day 3 each gravid female was separated in small cups lined with filter paper and filled with ¼ water into it. Each cup was covered with mosquito net and labelled. Cotton balls soaked with a sugar solution were kept on top of each cup. Females from five founder lines that laid a greater number of eggs and had 90-95% hatchability were selected. These selected lines were maintained separately to generate iso-female lines. The larvae and adults were given the same food as mentioned earlier. Emerged adults from each line were kept separately in mosquito rearing cages (Bugdorm-4S1515) (W17.5 x D17.5 x H17.5 cm) and allowed for sibling mating. On day 5, adult females of each line were blood fed and gravid females were separated in cups to obtain the next progeny in the similar way as mentioned above. The same procedure was continued till 5 generations and established a homozygous IndCh iso-female line.

### Photomap of polytene chromosome

Polytene chromosomes were prepared from the ovarian nurse cells collected from semi-gravid females of the IndCh iso-female line in the 40th generation as per the method of Ghosh & Shetty[39]. The semi-gravid females were anaesthetised and placed on a microslide in a drop of diluted Carnoy’s fixative solution (Carnoy’s fixative: distilled water, 1:19). The ovaries were pulled out and fixed in modified Carnoy’s fixative (methanol: acetic acid, 3:1) for 2-3 minutes. After fixation, the ovaries were stained with lacto acetic orcein for 15-20 minutes. After staining, 60% acetic acid was added and a clean coverslip was placed on the top of the stained sample. Gentle pressure was applied on the cover glass for squashing. The edges of the coverslip were sealed with nail polish. The slides were examined under microscope at 400X and 1000X, respectively for inversions. The inversion nomenclature and their frequency were recorded[5].

### Assembly

PacBio reads from the IndCh strain were assembled independently using Canu[20] and FALCON [23] assemblers. The resulting assemblies were combined using Quickmerge[21], where the Canu assembly was taken as reference and the FALCON assembly served as the query. Two rounds of Arrow polishing (PacBio gcpp v2.0.2) using PacBio reads, followed by two rounds of Pilon polishing (v1.22)[22] using 100x Illumina reads, were carried out on the merged assembly. HiC data from STE2 strain of *An. stephensi* was utilized to scaffold the contigs using SALSA[40, 41], where HiC reads were mapped to the contigs using the Arima Genomics mapping pipeline. DNA physical marker sequences for each chromosome arm published by Jiang *et al.*[6] were downloaded and aligned against the scaffolds using BLAST[42], in order to identify the chromosomal scaffolds. The scaffolds were stitched based on the order and orientation of the markers into pseudo-chromosomes. 100 N’s were added between two scaffolds during stitching. Physical marker sequences were realigned to the stitched chromosomes. Coordinates from the BLAST output, along with the stitched chromosome lengths, were utilized to generate a karyogram for visualization.

### Haplotype Phasing

The assemblers FALCON and FALCON-Unzip were used to phase the IndCh genome into haplotypes. Raw IndCh PacBio reads were used by the FALCON assembler to produce a set of primary contig files and an associate contig file representing the divergent allelic variants. Bubbles in the contig-assembly graph that result from structural variation between haplotypes are resolved as associate and primary contigs. The output of primary and associate contig files were then used by FALCON-Unzip to produce partially phased primary contigs (all_p_ctg.fa) and fully phased haplotigs (all_h_ctg.fa), which represent divergent haplotypes. After obtaining these results, phased polishing was performed by FALCON-Unzip using Arrow. This method of polishing preserves the haplotype differences by polishing the primary contigs and alternate haplotigs with reads that are binned into the two haplotypes. The final set of consensus primary and haplotig contigs were obtained named cns_p_ctg.fasta and cns_h_ctg.fasta respectively.

### Purge Haplotigs

The tool ‘Purge Haplotigs’[43] was used to determine the degree of heterozygosity in the IndCh strain. PacBio reads of IndCh strain were mapped to the consensus primary assembly (cns_p_ctg.fasta) obtained from FALCON-Unzip to obtain an aligned BAM file. Coverage analysis was performed on the BAM file to generate a read-depth histogram.

### Identification of 2Rb Breakpoints in UCI and IndCh

The tools HiC-Pro and Juicebox were used to generate a contact map for the genome of UCI (which is homozygous for the 2R*b* inverted form) by mapping the HiC reads from the STE2 strain (which is heterozygous for the 2R*b* inversion). The rationale was that the HiC reads corresponding to the standard form of 2R*b* will show up as an off-diagonal butterfly suggesting the position of breakpoints. Similarly, paired-end HiC reads from the UCI were mapped onto the IndCh assembly using HiC-Pro. As UCI is homozygous for the 2R*b* inversion, the HiC reads were expected to create a butterfly-like structure on the IndCh assembly, suggesting that it is the standard form with respect to the 2R*b* locus. These breakpoints were visualized and extrapolated from the contact map of the tool Juicebox. The breakpoints for the 2R*b* locus were deduced by mapping the PacBio and Illumina reads of IndCh on UCI genome to identify a gap, or lack of support for the reads for the inverted form. The same PacBio and Illumina reads from the IndCh strain were remapped on the IndCh assembly expecting that the standard form of 2R*b* locus in IndCh would not yield any gap near the coordinates obtained in the contact map. Alignment using ‘unimap’ was performed between the two assemblies of IndCh and UCI, to obtain the precise breakpoints for the 2R*b* inversion to consolidate with the findings from HiC-Pro.

### Repeat Analysis and Annotation

A database for the IndCh assembly was created using the ‘BuildDatabase’ utility. This database was submitted to the RepeatModeler version 2.0.1[44] along with the -LTRStruct parameter to predict the long terminal repeats. The custom library of consensus sequences <DATABASE_NAME>-families.fa from the RepeatModeler output and the -no_is parameter for skipping bacterial insertion elements were given as input for masking the repetitive elements using RepeatMasker version 4.1.0[45]

### Insect samples

In total, 82 mosquito samples which includes 4 lab strains TI Bangalore Lab (n=15), TII Chennai lab (n=15), TIII Delhi Lab (n=16), TIV Mangalore Lab (n=16) and 2 wild type strains from different geographical regions of India (Bangalore (n=10) and Mangalore (n=10)) were used in the present study. Lab strains were obtained courtesy of Dr. S. K. Ghosh (NIMR).

### Isolation of Genomic DNA

Briefly, the insect tissue was homogenized and the genomic DNA was isolated using Qiagen Genomic-tip (Qiagen). Later, fluorometric quantification of DNA was done using Qubit (Invitrogen). This procedure was adapted from our recent publication[3].

### Library preparation and sequencing

Whole Genome DNA libraries with average insert size of 200 bp were made using NEBNext® Ultra™ II DNA Library Prep Kit for Illumina® (New England Biolabs, 2016) using the protocol recommended by the company. Briefly, around 50 ng of DNA was used for library preparation, DNA was sheared using Adaptive Focused Acoustic technology (Covaris, Inc.) to generate fragments of length around 200 bp. The fragments were end repaired, 3′-adenylated, ligated with Illumina adapters, and PCR enriched with Illumina sequencing indexes. The size selection was performed using solid-phase reversible immobilization (SPRI) beads (Agencourt AMPure XP Beads) from Beckman Coulter. The quality and quantity of the libraries were evaluated using Qubit (Invitrogen) and TapeStation (Agilent). The libraries were diluted and pooled with an equimolar concentration of each library. Cluster generation was done using cBot (Illumina) and paired-end sequenced on Illumina HiSeq 2500 platform using TruSeq SBS Kit v3-HS (200 cycle) (Illumina, San Diego, CA) following the manufacturer’s recommendations. This procedure was adapted from our recent publication[3] .

### Gene Annotation

AUGUSTUS (version 3.2.3)[46], a eukaryotic gene prediction tool, was used to find protein-coding genes for the IndCh assembly and the UCI genome. AUGUSTUS uses ab initio gene prediction and reconciles the predicted gene structures with orthology to the proteome from a model organism. The model organism closest to *An. stephensi* was made available by AUGUSTUS for use in gene prediction as *Ae. aegypti*. AUGUSTUS provides a gff3 file delineating the exon–intron boundaries for each predicted gene along with its protein sequence. These gff files are uploaded to the browser, the link to which can be found in the section “Data Availability Section”. These gff files were used for gene liftover between UCI and IndCh. This procedure was adapted from our recent publication[3].

### Liftover

The two genomes were aligned using nucmer (NUCleotide MUMmer). Parameters ‘--maxmatch’ and ‘--noextend’ made sure to show all alignments regardless of their uniqueness, and to not extend clusters of alignments to make a longer alignment respectively so as to be able to detect structural variants. The delta file obtained from the nucmer run was converted to a text file containing coordinates of the alignment blocks between the two assemblies using the show-coords utility. This output file was converted to a bed file format, which contained the start and stop coordinate information of alignment blocks from the UCI genome with the corresponding chromosome, start, and stop information from the IndCh assembly of the alignment block was stored in the fourth column along with the strandedness. The bed file can be viewed on a genome browser along with the reference (UCI) to view genome liftover with a trackname titled ‘IndCh_UCI_BED’.

This genome liftover was then used as a base to lift the genes of the IndCh to the coordinates of UCI (from the gene annotation obtained from AUGUSTUS). For each gene in IndCh, an alignment block was chosen, which is from the same chromosome of UCI as the gene in IndCh, and which encompasses the start and stop codon of the gene/feature (exon or CDS). Based on the strandedness of the IndCh strain in the alignment block and the strandedness of the gene, the corresponding start and stop coordinates for the UCI genome were determined. This can be viewed on a genome browser as the track titled ‘IndCh_Gene’.

### Structural Variant Identification

PacBio structural variant calling and analysis tools, collectively called as PBSV (https://github.com/PacificBiosciences/pbsv) were used to discover structural variants in IndCh using UCI genome as reference. The PBSV pipeline has three main steps: 1) Alignment of IndCh PacBio reads to reference using ‘pbmm2 align’, 2) Discovery of structural variation signatures using ‘pbsv discover’ and 3) Calling of structural variants and assigning genotypes using ‘pbsv call’ options. The obtained VCF file from PBSV was used to analyse different structural variants like deletion, insertions, CNVs etc. from the ‘INFO’ column by filtering the ‘SVTYPE’.

### Structural Variant Annotation

SnpEff was used to annotate SVs and predict their effect on known genes. A snpEff database was created using the UCI reference genome and .gff file obtained from MAKER[9]. This was then used to annotate the SV file created using PBSV.

### PCR validation

Primers were designed to validate the 2R*b* breakpoints from the IndCh assembly using the tool ‘Primer-BLAST’. The forward primer for IndCh left breakpoint was (5’ GGGGATGGGAACGTGTTTCATA 3’), whereas the reverse primer sequence (5’ CATTCGCCACGTTTCAACTCAC 3’) was used to amplify the the region of IndCh chromosome 2R from 38,082,572 to 38,083,196 bp. Similarly, the forward primer for IndCh right breakpoint was (5’ TCGTCCCATTTCAGTCGGTAG 3’), whereas the reverse primer sequence (5’ ATGGTTGCTAAGCACGATGCAG 3’) was used to amplify the the region of IndCh chromosome 2R from 21,513,342 to 21,514,478 bp.

The DNA of individual mosquitoes were isolated from IndCh iso-female sibling colony and TII chennai lab colony using Qiagen Blood and Tissue Midi kit (Cat. No. 69506). The fluorometric quantification of DNA was done using NanoDrop (Thermo Scientific 2000 Spectrophotometers).

Each PCR reaction mixture (20 ul) contained 1 ul gDNA, 50 ng each forward and reverse primers (as mentioned above), 10 ul of Q5® Hot Start High-Fidelity 2X Master Mix and 8 ul of Nuclease free water added. A hot start of 95° C for 2 min. followed by 0.40 min denaturation, 70C annealing and 30 cycles of extension at 72° C. The PCR products were cleaned using a QIAquick PCR Purification Kit and given for Sanger sequencing.

### 2Rb Inversion Signature

Illumina reads for 15X coverage were generated for 82 samples, belonging to 2 wild and 4 lab populations. These reads were mapped to the high quality genome of UCI using bowtie2[47]. Variants were called using Samtools and the filtration was done upon the resulting VCF files using bcftools[48].

Fastq files of Illumina reads for a coverage of about 100X from UCI (SRR11672504) and STE2 (SRR1168951) were sourced from NCBI SRA (https://www.ncbi.nlm.nih.gov/sra) and mapped against the high-quality genome of UCI. The resulting VCF along with the VCF of the IndCh genome served as the training dataset for inversion signature. The UCI genome has a homozygous inverted 2R*b* genotype, STE2 is heterozygous for the 2R*b* genotype, and IndCh is homozygous to standard form of 2R*b* genotype as shown in the schematic 8, which shows the strategy for selecting the 22,650 signature SNPs, in that, these locations have homozygous alternate alleles in IndCh, heterozygous allele in STE2 and homozygous reference in UCI strain.

The alleles at the 22,650 positions were called from 82 individuals from 6 populations to attempt unsupervised clustering. The next step was to perform supervised clustering after cluster assignment for each of the 82 samples from the previous step to identify 3,781 most significant SNPs with a p-value of <0.0005. The number of SNPs were further reduced to 277 by selecting only those SNPs from the 3,781 which fall in clusters of more than 6 within a stretch of 1500 bases. Also, we identified 1740 SNPs within exon out of the 22,650 SNP to enable 2Rb genotyping directly from transcriptome data.

## ABBREVIATIONS

IndCh - India, Chennai strain; SNP - Single Nucleotide polymorphism; PCR - Polymerase Chain Reaction; LTR - Long Terminal Repeats; UCI - University of California, Irvine; HiC - High throughput chromatin conformation capture; NIMR - National Institute of Malaria Research; DNA - Deoxyribonucleic Acid; WHO - World Health Organization; RFLP - Restriction Fragment Length Polymorphism; BLAST - Basic Local Alignment Search Tool; SNV – Single Nucleotide Variant; CNV - Copy Number Variants; UTR - Untranslated Region; PCA - Principal Component Analysis; BAM - Binary Alignment Map; GFF - General Feature Format; CDS - Coding Sequences; PBSV - PacBio Structural Variant; VCF - Variant Call Format; GPS - Global positioning System; WGS - Whole Genome Sequencing.

## DATA AVAILABILITY AND MATERIALS

Assembly, raw PacBio/ Illumina reads used in the assembly and individual WGS dada used to validate 2R*b* signatures are uploaded to the NCBI server under the BioProject ID: PRJNA746765.

Download 22,650 SNPs, 3781 SNPs, and 1740 SNPs signatures at <u>Download</u> UCI Genome Browser- <u>JBrowse</u> (see Supplementary Table 2 for track information) IndCh Genome Browser - <u>JBrowse</u> (See Supplementary Table 3 for track information) As shown in the flowchart below, IndCh strain was derived from Chennai lab strain collected from an inner urban setting at a construction site at Annanagar, Chennai in 2016 (see Fig. 8 for GPS coordinates). The iso-female line was established to homogenize the population over 5 generations and 60 individual females were sent for sequencing. The iso-female line was continued until 20 generations and left to sustain as a separate lab colony for another 20 generations. All validation reported here including PCR and polytene chromosome work is done using the outgrown population and the original Chennai-lab strain.

For validation of segregating signatures for 2R*b* genotype WGS data were generated using the following lab-reared and wild populations.

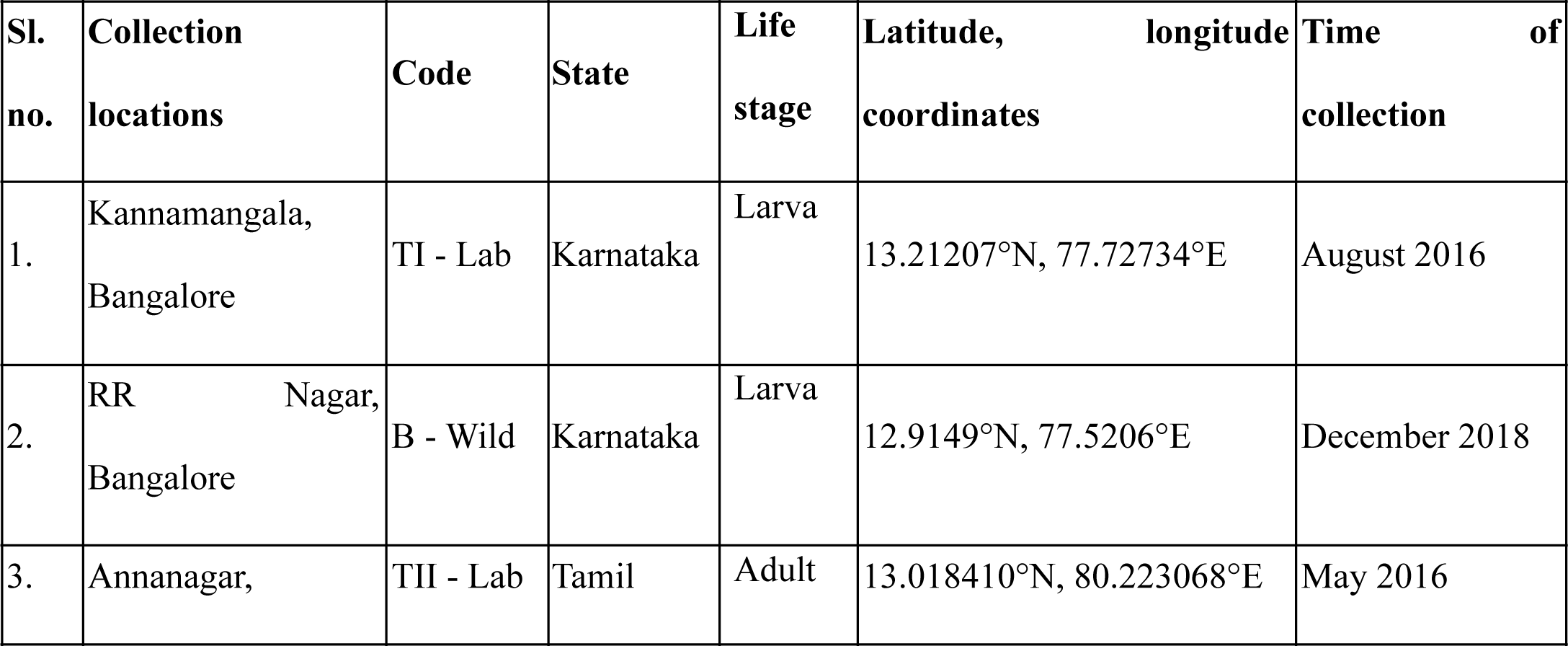
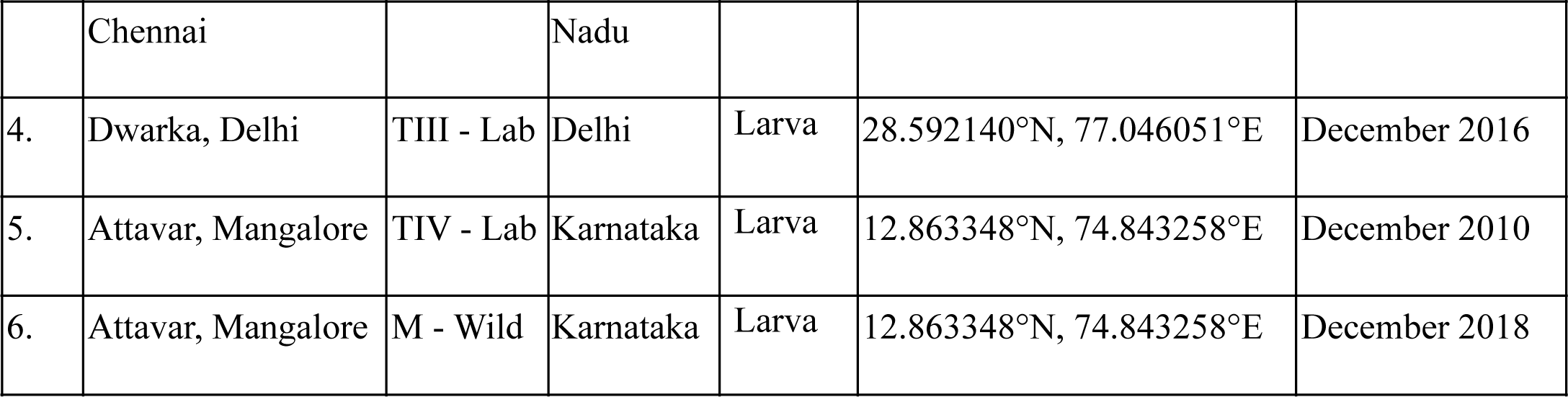

## Public Source of Data used in this report

Fastq files of Illumina reads for a coverage of about 100X from UCI (SRR11672504) and STE2 (SRR1168951) were sourced from NCBI SRA.

Developmental transcriptome data were obtained from the SRA database (<u>SRP013839</u>).

Scripts used for analysis can be obtained from <u>Github</u>. Additional Information related to genes of interest and other raw data can be obtained from the <u>IndCh website</u>.

## ACKNOWLEDGEMENT

The authors thank Nucleome Informatics Pvt. Ltd. for generating 60X Pacbio reads and 100X Illumina reads for IndCh. The authors thank Tata Institute for Genetics and Society (TIGS) India for funding researchers involved in this work and for funds for sequencing and Government of Karnataka for funding the computational infrastructure at IBAB.

## AUTHOR INFORMATION

Affiliations

Institute of Bioinformatics and Applied Biotechnology, Biotech Park, Electronic City Phase 1, Bengaluru 560100, India

Aditi Thakare, Tejashwini Alalamath, Himani Narang, Saurabh Whadgar, Kiran Paul, Shweta Shrotri, Raksha Rao, Bibha Choudhary and Subhasini Srinivasan.

Tata Institute for Genetics and Society, Center at inStem – GKVK campus, Bellary Road, Bengaluru 560065, India

Chitali Ghosh, Naveen Kumar, Sampath Kumar, Soumya M and Sunita Swain.

University of California, Irvine, CA 92697, USA

Mahul Chakraborty

University of California San Diego, La Jolla, CA 92093, USA

Suresh Subramani

National Institute of Malaria Research, Bangalore 562110, India

Susanta K. Ghosh

## Contributions

AT, writing the manuscript, for showing that IndCh is homozygous to standard form using bioinformatics, validating the breakpoints, comparative repeat analysis and primer designing; CG, for maintaining the chennai lab population, creating iso-female-line and validating 2Rb inversion using polytene chr; TA, extracting 2Rb signature and demonstrating that one-third of population is maintained in heterozygous form even in lab reared individuals, compiling refe; NK, Insectary work and PCR validation of sequence near 2Rb breakpoints in IndCh; HN, assembled the final genome; SW, identification of potential candidate genes within 2Rb locus and development of IndCh browser; KP, annotation of the genome, and analyzing the structural variants ; SShrotri, comparative 2Rb genomics and liftover of IndCh gene to UCI coordinates; SK, Insectary work and iso-female-line maintenance; SM, Insectary work and iso-female-line maintenance, RR, sequencing and library preparation of 92 individuals; MC, for initial quality check of raw data using FALCON assembly; BC, coordinated lab work at IBAB; SKG, collected, propagated and made available all the lab strains of An. stephensi; SureshS, designing the project and overall coordination; SunitaS, coordinating insectary work; SS, coordinating bioinformatics work and writing of the manuscript.

## REFERENCES

1. Sinka ME, Pironon S, Massey NC, Longbottom J, Hemingway J, Moyes CL, et al. A new malaria vector in Africa: Predicting the expansion range of *Anopheles stephensi* and identifying the urban populations at risk. Proc Natl Acad Sci U S A. 2020;117:24900–8. [DOI:10.1073/pnas.2003976117]

2. Sinka ME, Bangs MJ, Manguin S, Chareonviriyaphap T, Patil AP, Temperley WH, et al. The dominant *Anopheles* vectors of human malaria in the Asia-Pacific region: occurrence data, distribution maps and bionomic précis. Parasit Vectors. 2011;4:89. [DOI:10.1186/1756-3305-4-89]

3. Chida AR, Ravi S, Jayaprasad S, Paul K, Saha J, Suresh C, et al. A Near-Chromosome Level Genome Assembly of *Anopheles stephensi*. Front Genet. 2020;11:565626. [DOI:10.3389/fgene.2020.565626]

4. Kamali M, Sharakhova MV, Baricheva E, Karagodin D, Tu Z, Sharakhov IV. An integrated chromosome map of microsatellite markers and inversion breakpoints for an Asian malaria mosquito, *Anopheles stephensi*. J Hered. 2011;102:719–26. [DOI:10.1093/jhered/esr072]

5. Sharakhova MV, Xia A, Tu Z, Shouche YS, Unger MF, Sharakhov IV. A physical map for an Asian malaria mosquito, *Anopheles stephensi*. Am J Trop Med Hyg. 2010;83:1023–7. [DOI:10.4269/ajtmh.2010.10-0366]

6. Jiang X, Peery A, Hall AB, Sharma A, Chen X-G, Waterhouse RM, et al. Genome analysis of a major urban malaria vector mosquito, *Anopheles stephensi*. Genome Biol. 2014;15:459. [DOI:10.1186/s13059-014-0459-2]

7. Waterhouse RM, Aganezov S, Anselmetti Y, Lee J, Ruzzante L, Reijnders MJMF, et al. Evolutionary superscaffolding and chromosome anchoring to improve *Anopheles* genome assemblies. BMC Biol. 2020;18:1. [DOI:10.1186/s12915-019-0728-3]

8. Lukyanchikova V, Nuriddinov M, Belokopytova P, Liang J, Reijnders MJMF, Ruzzante L, et al. *Anopheles* mosquitoes revealed new principles of 3D genome organization in insects. bioRxiv. Cold Spring Harbor Laboratory; 2020;2020.05.26.114017. [DOI:10.1101/2020.05.26.114017]

9. Chakraborty M, Ramaiah A, Adolfi A, Halas P, Kaduskar B, Ngo LT, et al. Hidden genomic features of an invasive malaria vector, *Anopheles stephensi*, revealed by a chromosome-level genome assembly. BMC Biol. 2021;19:28. [DOI:10.1186/s12915-021-00963-z]

10. Coluzzi M, Petrarca V, di Deco MA. Chromosomal inversion intergradation and incipient speciation in Anopheles gambiae. Bollettino di zoologia. Taylor & Francis; 1985;52:45–63. [DOI:10.1080/11250008509440343]

11. Riehle MM, Bukhari T, Gneme A, Guelbeogo WM, Coulibaly B, Fofana A, et al. The Anopheles gambiae 2La chromosome inversion is associated with susceptibility to Plasmodium falciparum in Africa. Elife. 2017;6:e25813. [DOI:10.7554/eLife.25813]

12. Gray EM, Rocca KAC, Costantini C, Besansky NJ. Inversion 2La is associated with enhanced desiccation resistance in Anopheles gambiae. Malar J. 2009;8:215. [DOI:10.1186/1475-2875-8-215]

13. Coluzzi M. Inversion polymorphism and adult emergence in *Anopheles stephensi*. Science. 1972;176:59–60. [DOI:10.1126/science.176.4030.59]

14. Mahmood F, Sakai RK. Inversion polymorphisms in natural populations of *Anopheles stephensi*. Can J Genet Cytol. 1984;26:538–46. [DOI:10.1139/g84-086]

15. David J-P, Ismail HM, Chandor-Proust A, Paine MJI. Role of cytochrome P450s in insecticide resistance: impact on the control of mosquito-borne diseases and use of insecticides on Earth. Philos Trans R Soc Lond B Biol Sci. 2013;368:20120429. [DOI:10.1098/rstb.2012.0429]

16. Lobo NF, Sangaré DM, Regier AA, Reidenbach KR, Bretz DA, Sharakhova MV, et al. Breakpoint structure of the *Anopheles gambiae* 2Rb chromosomal inversion. Malar J. 2010;9:293. [DOI:10.1186/1475-2875-9-293]

17. Sharakhov IV, White BJ, Sharakhova MV, Kayondo J, Lobo NF, Santolamazza F, et al. Breakpoint structure reveals the unique origin of an interspecific chromosomal inversion (2La) in the Anopheles gambiae complex. Proc Natl Acad Sci U S A. 2006;103:6258–62. [DOI:10.1073/pnas.0509683103]

18. Cheng C, Tan JC, Hahn MW, Besansky NJ. Systems genetic analysis of inversion polymorphisms in the malaria mosquito *Anopheles gambiae*. Proc Natl Acad Sci U S A. 2018;115:E7005–14. [DOI:10.1073/pnas.1806760115]

19. Montanez-Gonzalez R, Pichler V, Calzetta M, Love RR, Vallera A, Schaecher L, et al. Highly specific PCR-RFLP assays for karyotyping the widespread 2Rb inversion in malaria vectors of the *Anopheles gambiae* complex. Parasit Vectors. 2020;13:16. [DOI:10.1186/s13071-019-3877-x]

20. Koren S, Walenz BP, Berlin K, Miller JR, Bergman NH, Phillippy AM. Canu: scalable and accurate long-read assembly via adaptive k-mer weighting and repeat separation. Genome Res. 2017;27:722–36. [DOI:10.1101/gr.215087.116]

21. Chakraborty M, Baldwin-Brown JG, Long AD, Emerson JJ. Contiguous and accurate de novo assembly of metazoan genomes with modest long read coverage. Nucleic Acids Res. 2016;44:e147. [DOI:10.1093/nar/gkw654]

22. Walker BJ, Abeel T, Shea T, Priest M, Abouelliel A, Sakthikumar S, et al. Pilon: an integrated tool for comprehensive microbial variant detection and genome assembly improvement. PLoS One. 2014;9:e112963. [DOI:10.1371/journal.pone.0112963]

23. Chin C-S, Peluso P, Sedlazeck FJ, Nattestad M, Concepcion GT, Clum A, et al. Phased diploid genome assembly with single-molecule real-time sequencing. Nat Methods. 2016;13:1050–4. [DOI:10.1038/nmeth.4035]

24. Mages CF, Wintsche A, Bernhart SH, Müller GA. The DREAM complex through its subunit Lin37 cooperates with Rb to initiate quiescence. Elife. 2017;6:e26876. [DOI:10.7554/eLife.26876]

25. Ylikallio E, Woldegebriel R, Tumiati M, Isohanni P, Ryan MM, Stark Z, et al. MCM3AP in recessive Charcot-Marie-Tooth neuropathy and mild intellectual disability. Brain. 2017;140:2093–103. [DOI:10.1093/brain/awx138]

26. Wickramasinghe VO, McMurtrie PIA, Mills AD, Takei Y, Penrhyn-Lowe S, Amagase Y, et al. mRNA export from mammalian cell nuclei is dependent on GANP. Curr Biol. 2010;20:25–31. [DOI:10.1016/j.cub.2009.10.078]

27. Falaleeva M, Pages A, Matuszek Z, Hidmi S, Agranat-Tamir L, Korotkov K, et al. Dual function of C/D box small nucleolar RNAs in rRNA modification and alternative pre-mRNA splicing. Proc Natl Acad Sci U S A. 2016;113:E1625–1634. [DOI:10.1073/pnas.1519292113]

28. Bard F, Casano L, Mallabiabarrena A, Wallace E, Saito K, Kitayama H, et al. Functional genomics reveals genes involved in protein secretion and Golgi organization. Nature. 2006;439:604–7. [DOI:10.1038/nature04377]

29. Neafsey DE, Waterhouse RM, Abai MR, Aganezov SS, Alekseyev MA, Allen JE, et al. Mosquito genomics. Highly evolvable malaria vectors: the genomes of 16 *Anopheles* mosquitoes. Science. 2015;347:1258522. [DOI:10.1126/science.1258522]

30. Shetty NJ, Hariprasad TPN, Sanil D, Zin T. Chromosomal inversions among insecticide-resistant strains of *Anopheles stephensi* Liston, a malaria mosquito. Parasitol Res. 2013;112:3851–7. [DOI:10.1007/s00436-013-3575-0]

31. Tubio JMC, Tojo M, Bassaganyas L, Escaramis G, Sharakhov IV, Sharakhova MV, et al. Evolutionary dynamics of the Ty3/gypsy LTR retrotransposons in the genome of *Anopheles gambiae*. PLoS One. 2011;6:e16328. [DOI:10.1371/journal.pone.0016328]

32. Chakraborty M, Emerson JJ, Macdonald SJ, Long AD. Structural variants exhibit widespread allelic heterogeneity and shape variation in complex traits. Nat Commun. 2019;10:4872. [DOI:10.1038/s41467-019-12884-1]

33. Carballar-Lejarazú R, Jasinskiene N, James AA. Exogenous gypsy insulator sequences modulate transgene expression in the malaria vector mosquito, *Anopheles stephensi*. Proc Natl Acad Sci U S A. 2013;110:7176–81. [DOI:10.1073/pnas.1304722110]

34. Love RR, Redmond SN, Pombi M, Caputo B, Petrarca V, Della Torre A, et al. In Silico Karyotyping of Chromosomally Polymorphic Malaria Mosquitoes in the *Anopheles gambiae* Complex. G3 (Bethesda). 2019;9:3249–62. [DOI:10.1534/g3.119.400445]

35. Ayala D, Ullastres A, González J. Adaptation through chromosomal inversions in *Anopheles*. Front Genet. 2014;5:129. [DOI:10.3389/fgene.2014.00129]

36. Ghosh SK, Ghosh C. New Ways to Tackle Malaria [Internet]. Vector-Borne Diseases - Recent Developments in Epidemiology and Control. IntechOpen; 2019 [cited 2021 Aug 11]. Available from: https://www.intechopen.com/chapters/69269

37. Gunathilaka N, Ranathunge T, Udayanga L, Abeyewickreme W. Efficacy of Blood Sources and Artificial Blood Feeding Methods in Rearing of *Aedes aegypti* (Diptera: Culicidae) for Sterile Insect Technique and Incompatible Insect Technique Approaches in Sri Lanka. Biomed Res Int. 2017;2017:3196924. [DOI:10.1155/2017/3196924]

38. Ghosh C & Shetty NJ. Mode of inheritance of fenitrothion resistance in *Anopheles stephensi*. J Cytol Genet. 1999;34:141–6.

39. Chaitali Ghosh, N J Shetty. Tests for association of fenitrothion resistance with inversion polymorphism in the malaria vector, Anopheles stephensi Liston. Nucleus-Calcutta-International Journal of Cytology. 2004;47:164–8.

40. Ghurye J, Rhie A, Walenz BP, Schmitt A, Selvaraj S, Pop M, et al. Integrating Hi-C links with assembly graphs for chromosome-scale assembly. PLoS Comput Biol. 2019;15:e1007273. [DOI:10.1371/journal.pcbi.1007273]

41. Ghurye J, Pop M, Koren S, Bickhart D, Chin C-S. Scaffolding of long read assemblies using long range contact information. BMC Genomics. 2017;18:527. [DOI:10.1186/s12864-017-3879-z]

42. Altschul SF, Gish W, Miller W, Myers EW, Lipman DJ. Basic local alignment search tool. Journal of Molecular Biology. 1990;215:403–10. [DOI:10.1016/S0022-2836(05)80360-2]

43. Roach MJ, Schmidt SA, Borneman AR. Purge Haplotigs: allelic contig reassignment for third-gen diploid genome assemblies. BMC Bioinformatics. 2018;19:460. [DOI:10.1186/s12859-018-2485-7]

44. Flynn JM, Hubley R, Goubert C, Rosen J, Clark AG, Feschotte C, et al. RepeatModeler2 for automated genomic discovery of transposable element families. Proc Natl Acad Sci USA. 2020;117:9451–7. [DOI:10.1073/pnas.1921046117]

45. Smit, AFA, Hubley, R. RepeatModeler Open-1.0. [Internet]. 2008 [cited 2021 Aug 11]. Available from: <http://www.repeatmasker.org>

46. Hoff KJ, Stanke M. Predicting Genes in Single Genomes with AUGUSTUS. Curr Protoc Bioinformatics. 2019;65:e57. [DOI:10.1002/cpbi.57]

47. Langmead B, Salzberg SL. Fast gapped-read alignment with Bowtie 2. Nat Methods. 2012;9:357–9. [DOI:10.1038/nmeth.1923]

48. Danecek P, Bonfield JK, Liddle J, Marshall J, Ohan V, Pollard MO, et al. Twelve years of SAMtools and BCFtools. Gigascience. 2021;10:giab008. [DOI:10.1093/gigascience/giab008]

