## Supplementary for "The genome trilogy of *Anopheles stephensi*, an urban malaria vector, reveals structure of a locus associated with adaptation to environmental heterogeneity"

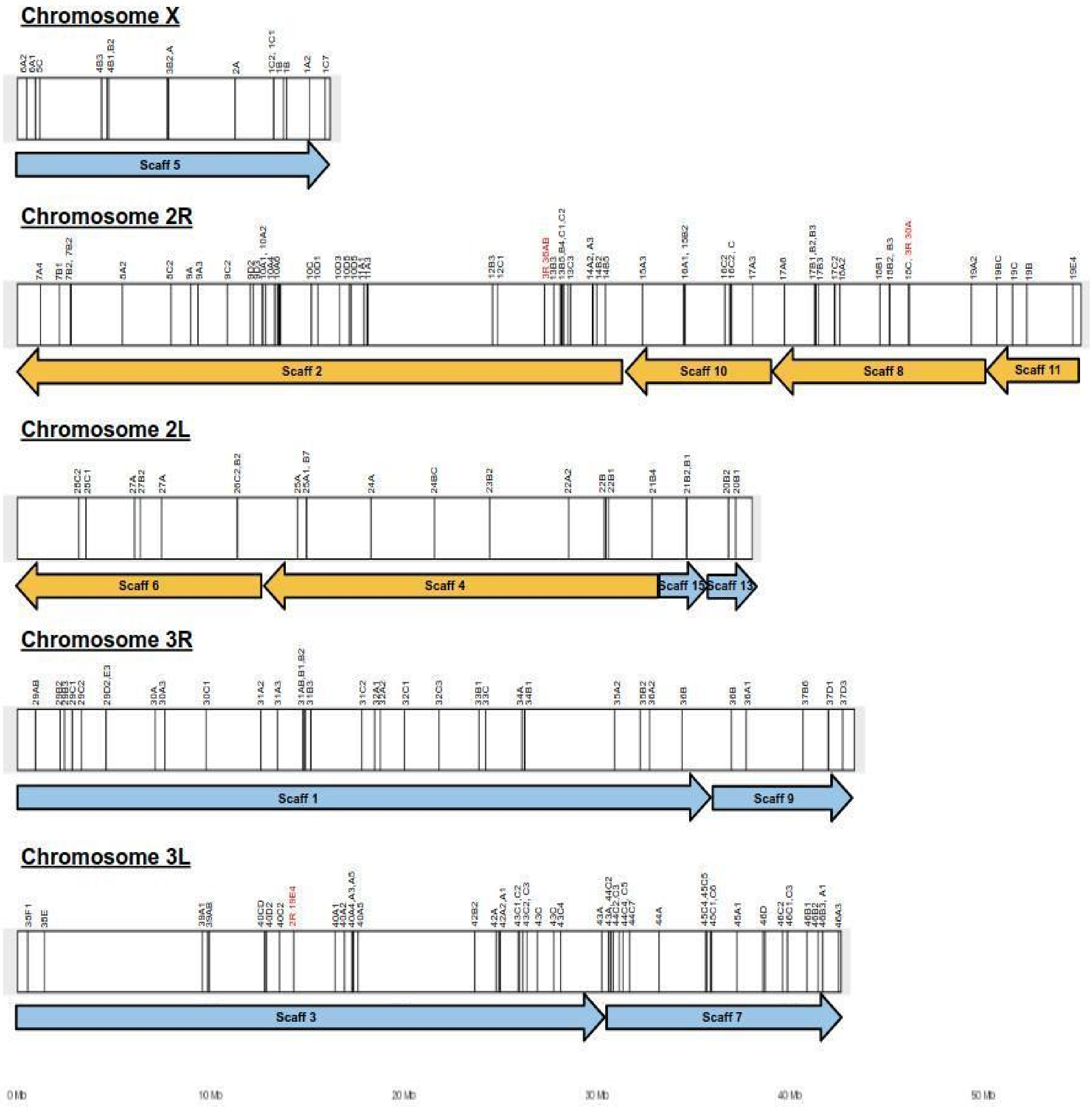


*Supplementary Fig. 1: Karyogram of scaffolds from HiC-based assembly stitched using physical markers for IndCh.*

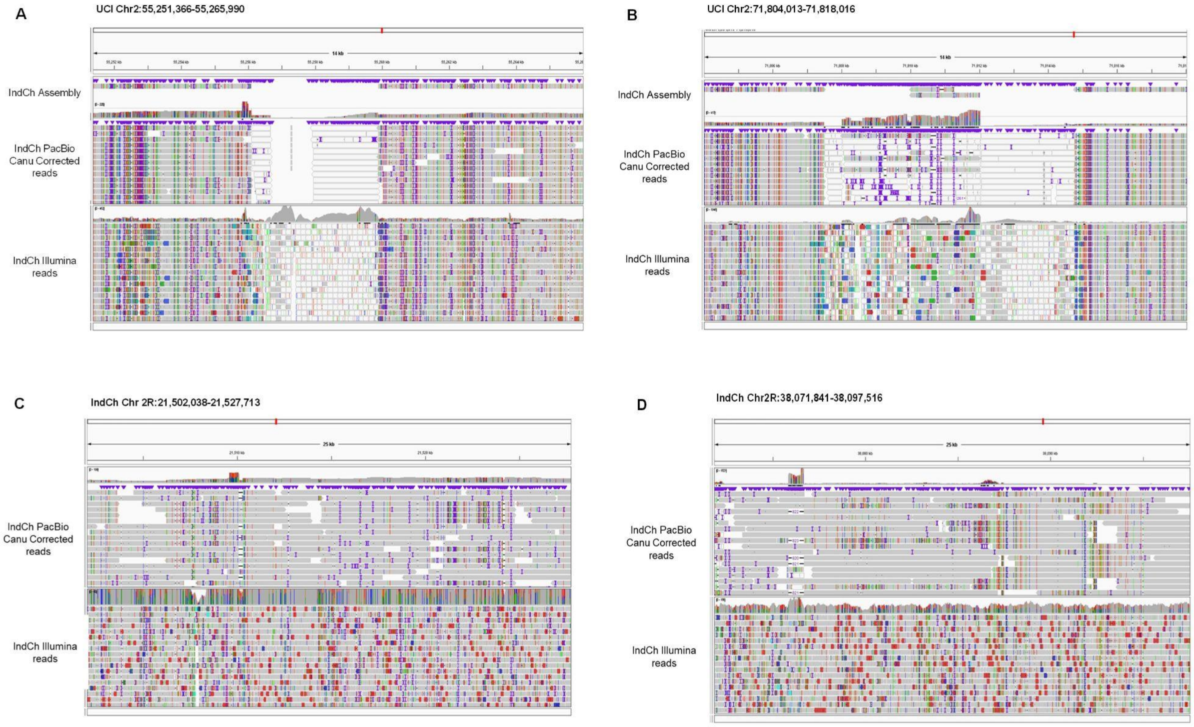


*Supplementary Fig. 2: Integrated Genome Viewer (IGV) visualization of CANU corrected IndCh PacBio reads (top) and IndCh Illumina reads (bottom) mapped on (A) left breakpoint of UCI assembly (B) right breakpoint of UCI Assembly (C) left breakpoint of IndCh assembly (D) right breakpoint of IndCh assembly.*

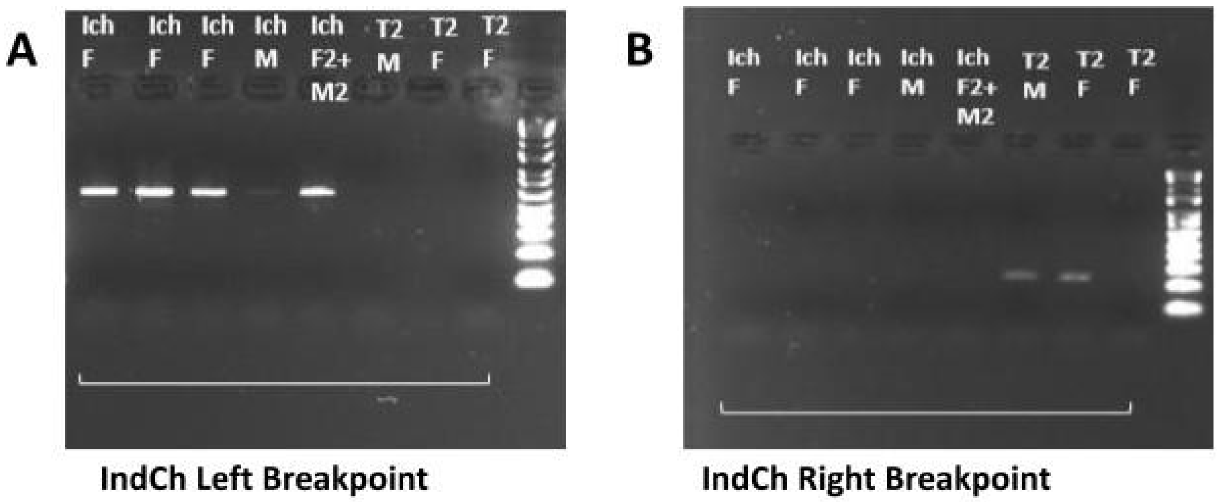


*Supplementary Fig. 3: (A) The left breakpoint from the IndCh assembly amplified in outgrown IndCh (Ich) population and from original Chennai lab population (T2) from which IndCh strain was derived (B) The right breakpoint from the IndCh assembly amplified only from Chennai lab population (T2). F is 'Individual Female Mosquito', M is 'Individual Male Mosquito', F2+M2 is two female mosquitoes and two male mosquitoes.*

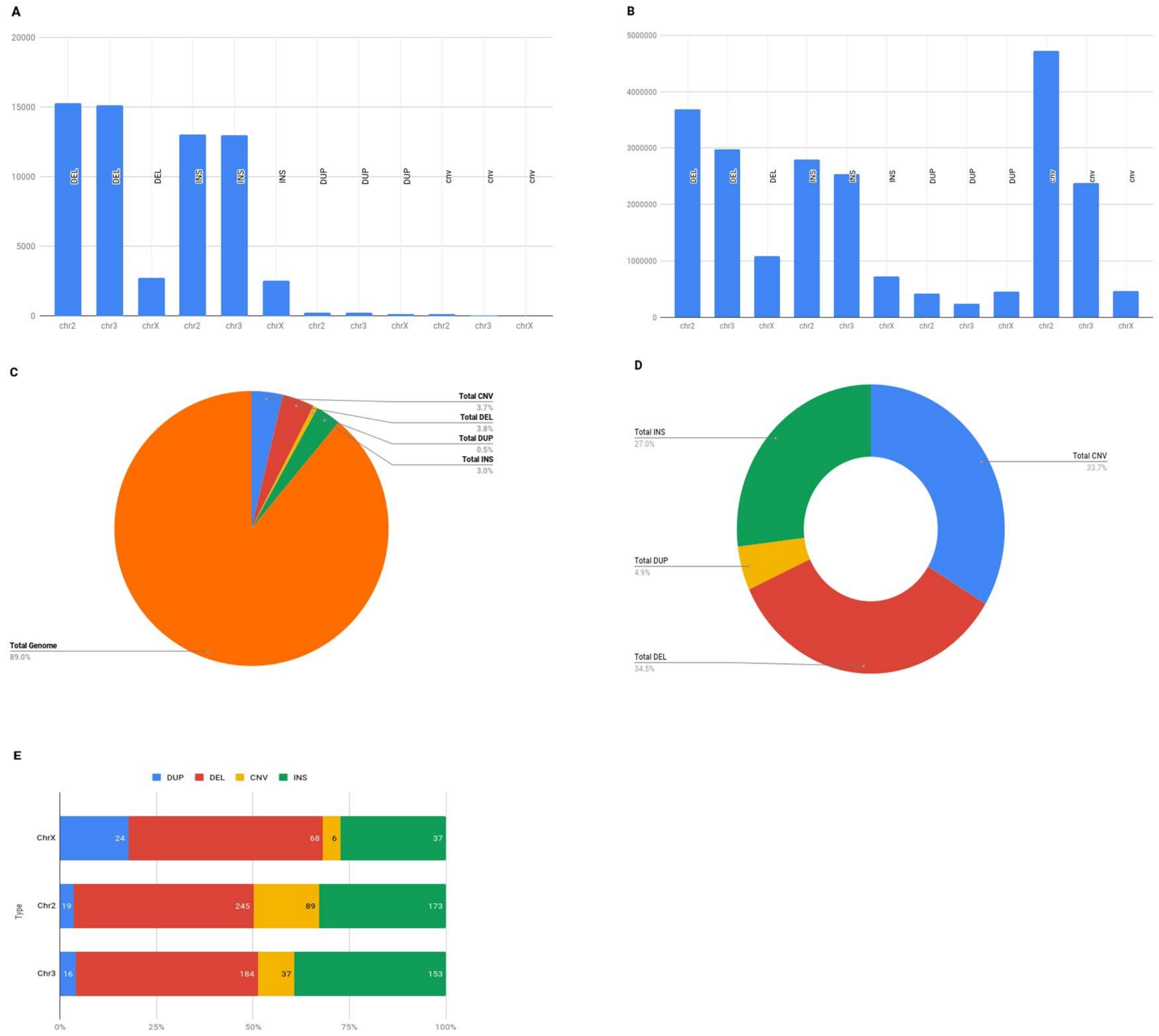


*Supplementary Fig. 4: (A) Shows number of structural variants present per chromosome, (B) shows the number of base pairs involved in each type of SVs per chromosome, (C) shows percent base pairs from the whole genome (chr2,3 and X) implicated in SVs. CNV and DEL accounts for almost 6% of the genome, (D) shows relative abundance of SV, DEL, INS, DUP and CNV, (E) shows relative distribution of SVs higher than 5 Kbp length.*

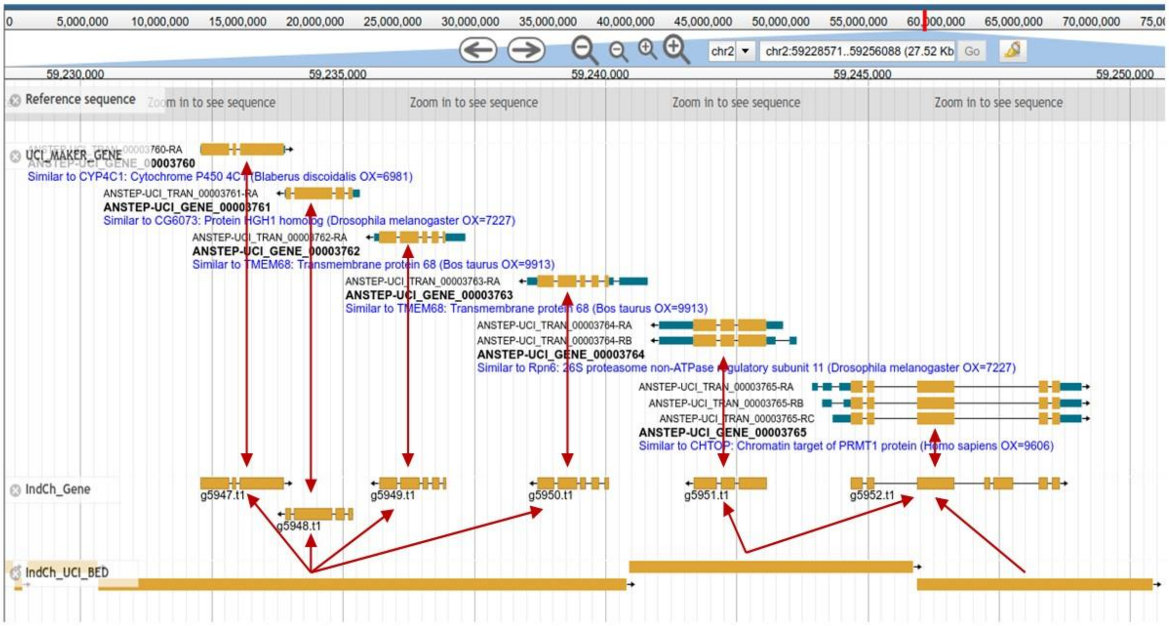


*Supplementary Fig. 5: A snapshot of the genome browser showing genome and gene liftover. The first track, "Reference sequence", is the UCI sequence. The next track, "UCI_MAKER_GENE", shows the Maker predicted genes for the UCI strain. The track "IndCh_Gene" shows the IndCh Augustus predicted genes lifted over to UCI coordinates. The track "IndCh_UCI_BED" shows the alignment blocks between IndCh and UCI on UCI coordinates. The six UCI genes shown here within the locus chr2:59,228,566-59,249,465 of UCI genome have corresponding genes in IndCh (marked with double headed arrows). Four alignment blocks used to lift over the genes are also shown.*

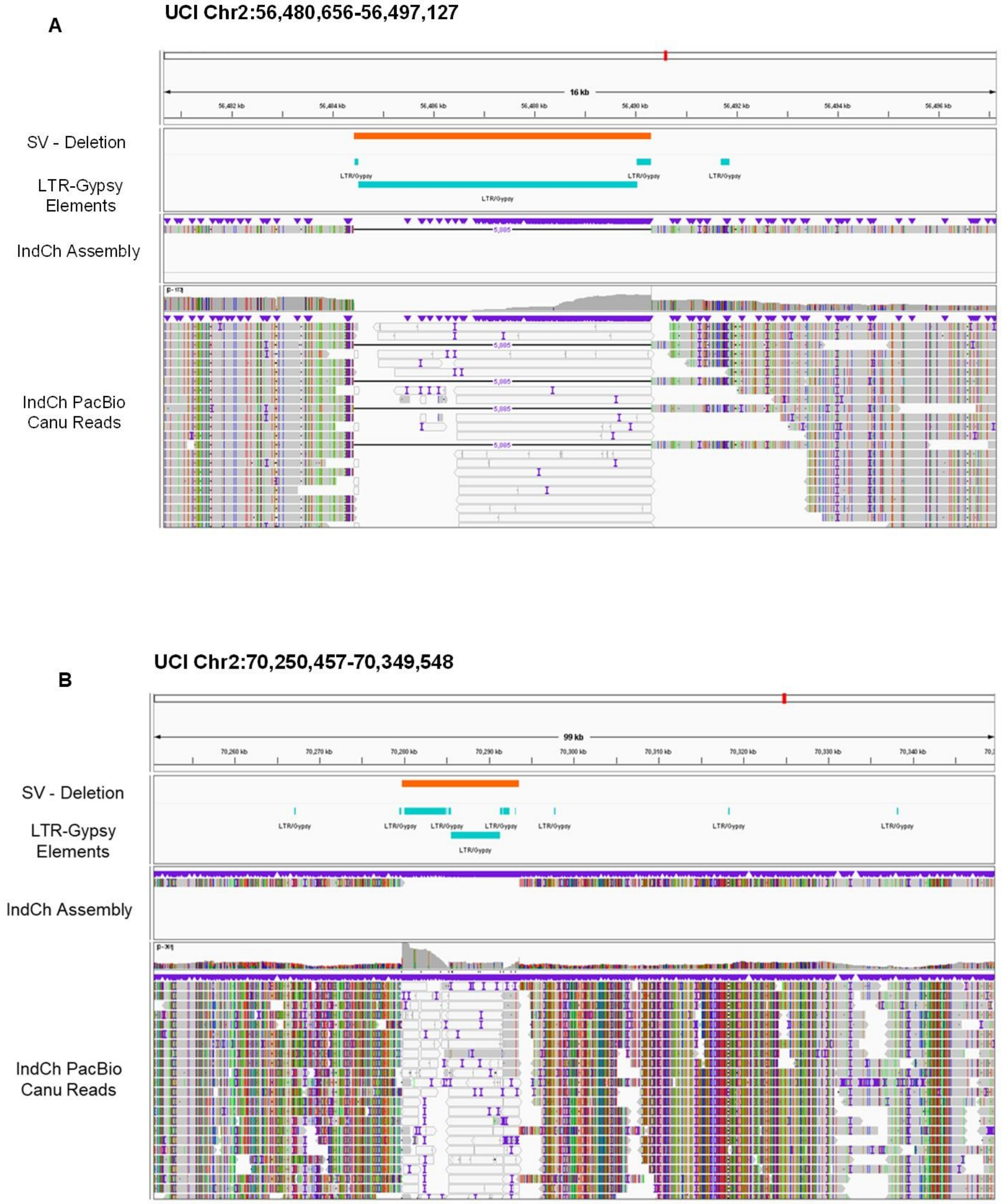


*Supplementary Fig. 6: Two examples (A and B) of IGV visualization supporting two LTR-Gypsy elements deleted from the IndCh assembly (top) and IndCh PacBio reads (bottom) when compared against the UCI genome.*

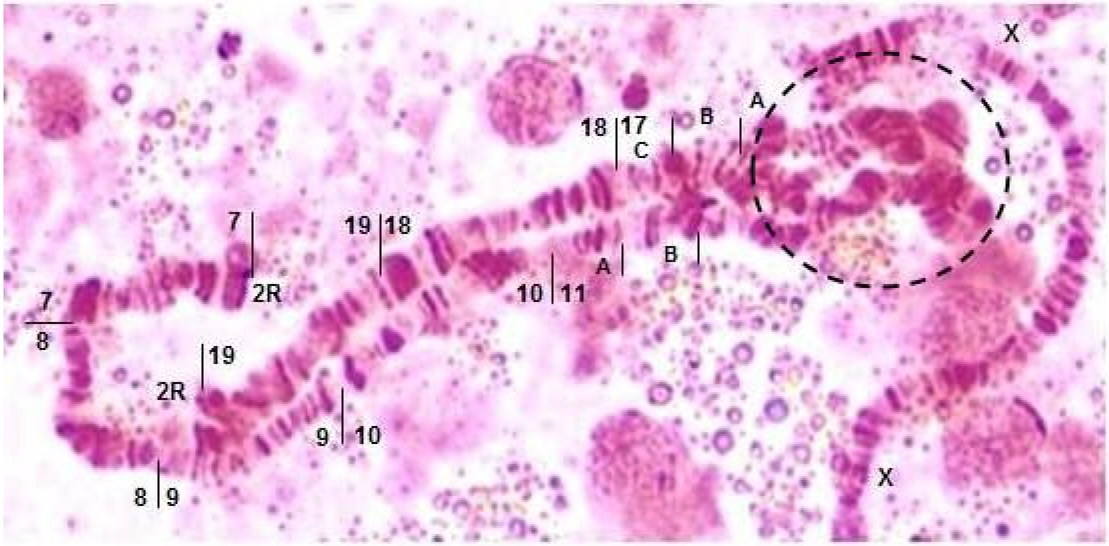


*Supplementary Fig. 7: Photomap supporting heterozygous 2Rb inversion in an individual. The black dotted circle shows the loop formation exhibiting the heterozygous form in 3.18% of the IndCh outgrown iso-female line.*

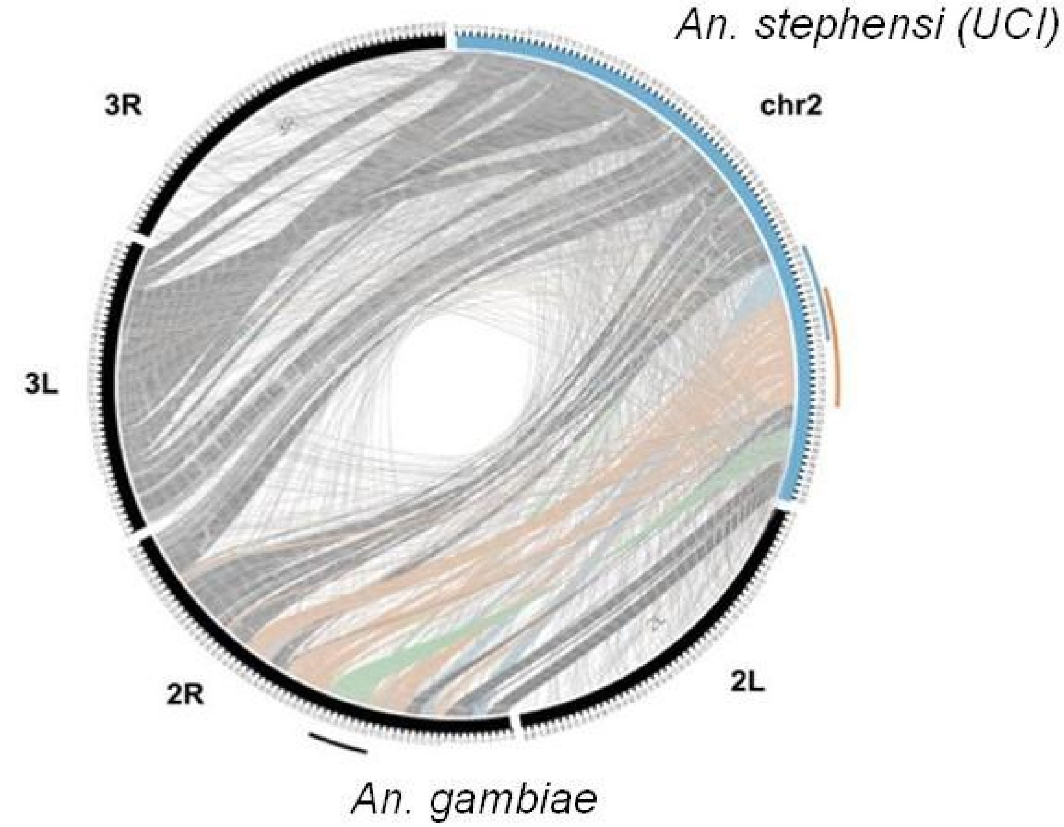


*Supplementary Fig. 8: Comparison between An.stephensi and An.gambiae genomes, showing no synteny within the 2Rb locus. Blue arc represents the 2Rb inversion locus in An. stephensi, orange arc is for the 2Ri inversion An. stephensi. Black arc shows the 2Rb inversion of An. gambiae. The colored connections highlight synteny.*

| From PBSV | Chr X | Chr 2 | 2R*b* | Chr 3 | Genome-wide | Genome-wide validation from PBSV with SVMU |
|---|---|---|---|---|---|---|
| No. of deletions/insertions (>= 5 Kbp) in IndCh | 63/20 | 244/151 | 56/28 | 183/101 | 490/272 | 445/248 |
| No. of deletions/insertions (>= 5 Kbp) in IndCh intersecting with the LTR-Gypsy elements (>= 1 Kbp) | 32/12 | 139/58 | 31/10 | 99/31 | 270/101 | 249/94 |
| No. of LTR-Gypsy (> 1 Kbp) elements spanning the deletions/insertions in IndCh | 47/12 | 162/43 | 34/11 | 117/28 | 326/83 | 297/105 |
| No. of the intersected LTR-Gypsy elements spanning in UCI genes (purple) / IndCh genes (brown) | 36/6 | 159/25 | 38/5 | 105/19 | 300/50 | NA |
| No. of reported LTR-Gypsy elements in UCI (>= 1 Kbp) | 413 | 347 | 50 | 209 | 969 | NA |
| No. of predicted LTR-Gypsy elements in IndCh (>= 1 Kbp) | 24 | 112 | 24 | 93 | 229 | NA |

*Supplementary Table 1: Stats showing deletions and insertions from PBSV of more than 5 Kbp in length observed in IndCh and their intersection with the LTR-Gypsy elements of length more than 1 Kbp. The last column validates the deletions and insertions from PBSV with SVMU.*

| No. | File Type | Track Name | Track Content |
|---|---|---|---|
| 1 | FASTA | Reference Sequence | *Anopheles stephensi genome 2.0 (UCI)* |
| | | Augustus Gene | Augustus predicted gene for UCI reference genome |
| | | IndCh Gene | IndCh-UCI Lifted Over gene |
| 2 | GFF | UCI Maker Gene | UCI MAKER Gene |
| | | IndCh_UCI_Illumina_Bowtie2 | IndCh Illumina reads mapped on UCI using Bowtie2 |
| | | IndCh_UCI_Pacbio_CANU | IndCh Pacbio CANU corrected read mapped on UCI |
| 3 | BAM | UCIwg_IndchNI | IndCh whole genome mapped on UCI using unimap |
| | | IndCh_PB_UCI-BND | BND type structural variant predicted by PBSV |
| | | IndCh_PB_UCI-CNV | Copy Number Variant structural variant predicted by PBSV |
| | | IndCh_PB_UCI-DEL | Deletion structural variant predicted by PBSV |
| | | IndCh_PB_UCI-INS | Insertion type structural variant predicted by PBSV |
| | | IndCh_PB_UCI-INS_DUP | Insertion-Duplication type structural variant predicted by PBSV |
| | | IndCh_PB_UCI-INV | Inversion structural variant predicted by PBSV |
| | | IndCh_PB_UCI-SPLIT_DUP | Split-Duplication type structural variant predicted by PBSV |
| | | UCI_PBSV | Structural variant predicted by PBSV with IndCh PacBio reads mapped on UCI reference |
| 4 | VCF | SVMU_SV | Structural variant predicted by SVMU |
| | | IndCh_UCI_BED | IndCh mapped on UCI |
| | | UCI_Repeats_their | Repeats predicted for UCI from PMID: 33568145 |
| | | R222-A01_aligned | Males (5-7 days old) |
| | | R222-B01_aligned | Females (5-7 days old ) unfed |
| | | R222-C01_aligned | Females (3-6 hrs Post Blood Meal) |
| | | R222-D01_aligned | Females (24 hrs Post Blood Meal) |
| | IsoSeq | R223-A01_aligned | Females (48 hrs Post Blood Meal) |

6 BAM

| | | | |
|---|---|---|---|
| | | R223-B01_aligned | Females (72 hrs Post Blood Meal) |
| 8 | BED | Physical_Marker | BLAST Physical Marker for genome |

*Supplementary Table 2 : Track details of JBrowse for UCI genome.*

| **No.** | **File Type** | **Track Name** | **Track Content** |
|---|---|---|---|
| 1 | FASTA | Reference sequence | IndCh genome as reference |
| 2 | GFF | Augustus_Gene | Genes predicted for IndCh genome using the 'Augustus' tool |
| | | Physical_Marker | BLAST of physical markers on the IndCh assembly |
| 3 | BED | 2RIIndchNI_REP | Repeat Elements predicted for IndCh |
| | | IndCh2R_UCI2 | Chromosome 2 of UCI mapped on IndCh Chr 2R |
| | | IndChNI_Illumina_Bowtie | IndCh Illumina reads mapped on IndCh genome using 'bowtie2' |
| 4 | BAM | IndCh_NI_PacBioCANU | IndCh PacBio Canu corrected reads mapped on IndCh genome using 'minimap2' |

*Supplementary Table 3 : Track details of JBrowse for IndCh genome.*

**Sequence spanning IndCh chromosome 2R left breakpoint**

CTTTGAAATGAATTAGATGATTTAAATCATCACGCAGCGTTATTTCAGCTCACTTTTGA
TTTAATTCGCCATGCCGATTAACCACTCACTCACTACTTTTTAACTCACTCACTATGAA
TGGCTCTCTC

*Supplementary Text 1 : Highlighted in blue are the 78 bps in IndCh near the 2Rb left breakpoint.*

**Sequence spanning IndCh chromosome 2R right breakpoint**

ACACGGGTAAGAATATAATCCAATCATATCTTTAGCGAGCTGCAAATTAGTGATTTTC

*Supplementary Text 2 : Highlighted in pink are the 8 bps in IndCh near the 2Rb right breakpoint.*
